# Unmasking Hidden Systemic Effects of Neurodegenerative Diseases: A Two-Pronged Approach to Biomarker Discovery

**DOI:** 10.1101/2023.11.23.568435

**Authors:** Sandra I. Anjo, Miguel Rosado, Inês Baldeiras, Andreia Gomes, Diana Pires, Cátia Santa, Joana Pinto, Cristina Januário, Isabel Santana, Ana Verdelho, Alexandre de Mendonça, Miguel Castelo-Branco, Bruno Manadas

## Abstract

Identification of reliable blood biomarkers for neurodegenerative diseases (NDs) is crucial for translational and clinical research. However, conventional omics struggle with blood samples complexity, hindering desired outcomes. In this work the potential of High Molecular Weight (HMW) fractionation under non-denaturing conditions as a complementary approach to the conventional proteomics for identifying serum biomarkers in NDs was explored. A cohort of 58 serum samples of Alzheimer’s disease (AD), Parkinson’s disease (PD) patients and control (CT) individuals was used to compare the two proteomics strategies: i) direct analysis of whole serum and ii) non-denaturing fractionation using 300 kDa cut-off filters (HMW serum).

Although both approaches quantified a similar set of proteins, each approach captured a distinct subset of differentially altered proteins, suggesting that HMW fractionation identified additional types of alterations beyond conventional protein level changes. A discriminant model combining altered proteins from both datasets effectively distinguished between the three groups (AUC = 0.999 and median sensitivity and specificity of 97.4% and 91.7%, respectively). Importantly, this performance surpassed that of any model created using each method individually.

Altogether, this work demonstrated that HMW fractionation can be a valuable complementary method to direct serum analysis and could enhance biomarker discovery. The 10 proteins included in the model (5 from each strategy), comprise clear evidence for the contribution of apolipoproteins for the diagnosis of NDs, revealing potential changes within lipid metabolism and the organization of macromolecules and their complexes, thereby uncovering effects that remain hidden from a conventional serum proteome analysis.

## Introduction

Proteins can be directly or indirectly related to a myriad of diseases, thereby being important targets of biomarker research, which remains a pivotal aspect of clinical research. However, biomarker discovery based on conventional proteomics strategies has not yet yielded substantial practical applications. Blood and its fractions are the most studied samples for biomarker identification, not only due to easy accessibility but also because of its interactions with most tissues in the body, potentially reflecting disease-related alterations [1, 2]. However, both plasma and serum proteomes have a large dynamic range of protein concentrations [1], with the detection of those least abundant being masked by the most abundant. Despite all the technical developments in proteomics quantification by mass spectrometry (MS) [3], the high dynamic range still precludes the complete characterization of these samples, which may be one of the reasons why the identification of relevant biomarkers in plasma/serum by MS remains challenging [4]. Therefore, it is essential to have alternative approaches to the direct analysis of these highly demanding biofluids. Several approaches have been developed to reduce sample complexity [5], such as size exclusion chromatography and electrophoretic separation methods. When dealing with numerous samples, a more straightforward approach that could serve as an alternative to direct sample analysis is centrifugal ultrafiltration [5]. This approach relies on the separation of sample components based on size exclusion, which is correlated to molecular weight (MW). This method has typically been used to study the low molecular weight proteome of serum and plasma samples, using filters between 20 kDa and 40 kDa [6, 7], but it may also be used to study other proteome fractions. Alzheimer’s Disease (AD) and Parkinson’s Disease (PD) are inherently linked to protein aggregation [8] and, therefore, to the formation of high molecular weight (HMW) protein complexes. Fractionation of peripheral fluid samples from patients with these diseases, focused on the HMW proteome, could be useful to eliminate high abundance proteins and to study protein aggregates that may be present in circulation. In this study, we propose using centrifugal ultrafiltration focused on the HMW proteome, not as an alternative but as a complementary analysis to the investigation of unfractionated samples, to potentially assess the restructuring of molecular complexes as a distinctive feature of disease. To evaluate our hypothesis, the same set of samples was subjected to the proteomics analysis of the whole serum and fractionation using centrifugal ultrafiltration with 300 kDa molecular weight cut-off (MWCO) filters in a non-denaturing environment (henceforth referred to as HMW fractionation). The data obtained from both approaches were directly compared and merged to identify potential biomarkers. This study used a cohort consisting of patients with neurodegenerative disorders, specifically Alzheimer’s and Parkinson’s disease patients, and healthy controls (CT) to test the significance of using HMW fractionation as a complementary tool for biomarker discovery. These disorders were chosen as they are considered as potential proteinopathies, making them promising targets for this research.

## Materials and Methods

### Participants

A total of 58 serum samples were used in this study, comprising 3 groups of individuals: AD (n = 22), PD (n = 24), and CT (n = 12). The PD patients were recruited at the Movement Disorders Units of the Neurological Department of the CHUC, where they were assessed by a movement disorders specialist and were diagnosed according to the criteria defined by the UK Parkinsons’s Disease Society Brain Bank [9]. The exclusion criteria for these patients consisted of severe dementia (as indicated by a Mini-Mental State Exam score below 15), any psychiatric disorder, or other forms of Parkinsonism. The clinical group of individuals with AD diagnosis was recruited and prospectively evaluated by two experienced neurologists at Memoclínica and the Neurology Department of the CHUC. The standard criteria for the diagnosis of AD were the Diagnostic and Statistical Manual of Mental Disorders—fourth edition (DSM-IV-TR) [10] and the National Institute on Aging and the Alzheimer’s Association Workgroup [11]. To ensure the homogeneity of the sample, only patients who met the following criteria were included: they were in a stable condition, did not sustain recent changes in medication, and did not have ophthalmological or neurological/psychiatric conditions other than AD. The CT group was composed of age- and gender-matched individuals from the community with no history of cognitive deterioration, neurological or acquired central nervous system (CNS) disorders, traumatic brain injury, or psychiatric disorders. The CT group was also submitted to a brief cognitive assessment to exclude the presence of cognitive impairment.

### Serum processing for proteomics analysis

Two different strategies were used to obtain a more comprehensive proteomic characterization of serum samples, namely: i) direct analysis of whole serum and ii) HMW fractionation through ultrafiltration using 300 kDa cut-off filters (HMW serum).

For each sample, 5 µL were used for direct serum analysis and 82.5 µL for HMW serum fractionation. Additionally, three sample pools were prepared by combining aliquots of all the samples, for the AD pool, PD pool, and CT pool, respectively. Pooled samples were used for Data-dependent acquisition (DDA) experiments to build a specific protein library to be used in data-independent acquisition (DIA) analysis and were subjected to the same sample processing as the individual samples. Before processing, all samples were spiked with the same amount of an internal standard (IS) to account for sample loss [12]. Different internal standards were used depending on the type of analysis: MBP-GFP [12] in the case of the whole serum approach, while equine ferritin, commonly available as one of the standards in the Gel Filtration Calibrants Kit for High Molecular Weight proteins (GE28-4038-42), for the HMW fractionation approach.

For the direct analysis of whole serum, the samples were diluted in Laemmli buffer, followed by denaturation for 5 min at 95°C and cysteine alkylation with acrylamide, and the total volume in all samples was subjected to in-gel digestion using the Short-GeLC for subsequent quantitative analysis by LC-MS/MS-DIA [13].

Samples subjected to HMW fractionation were ultrafiltrated using 300 kDa cut-off filters (Vivaspin^®^ 500 Polyethersulfone, 300 kDa (Sartorius)) pre-conditioned to PBS. Serum samples were diluted into 200 µL of PBS and subjected to 20 min centrifugation at 14,500× g at 4 °C followed by an additional washing step with another 200 µL of PBS. In some cases, the washing step was repeated until the retentate volume did not exceed 50 µL. The resulting retentates, the HMW fraction, were collected into a new LoBind^®^ microcentrifuge tube and precipitated with ice-cold acetone [14]. The precipitated pellets were resuspended into 30 µL of a solution containing 2% SDS (v/v) and 1 M of Urea, always aided by sonication (VibraCell 750 watt-Sonics ^®^) with ice in the cup horn (2 min. pulse duration, at 1 second intervals, and with 40% amplitude). Afterward, concentrated Laemmli Buffer was added to the samples, followed by a 30 min incubation to reduce the samples and a 20 min incubation with iodoacetamide for cysteine alkylation [15]. The total volume in all samples was subjected to in-gel digestion as previously specified [13].

### Mass spectrometry data acquisition

Samples were analyzed on a NanoLC™ 425 System (Eksigent^®^) couple to a TripleTOF™ 6600 System (Sciex^®^) using DDA for each fraction of the pooled samples for protein identification and SWATH-MS acquisition of each sample for protein quantification. Peptides were resolved by micro-flow liquid chromatography on a MicroLC column ChromXP™ C18CL (300 µm ID × 15 cm length, 3 µm particles, 120 Å pore size, Eksigent^®^) at 5 µL/min. The liquid chromatography program was performed as follows with a multistep gradient: 2 % to 5 % mobile phase B (0-2 min), 5 % to 28 % B (2-50 min), 28% to 35% B (50-51 min), 35 to 98% of B (50–52 min), 98% of B (52-61 min), 98 to 2% of B (61–62 min), 2% of B (68 min). Mobile phase A was composed of 0.1 % formic acid (FA) with 5% dimethyl sulfoxide (DMSO), and mobile phase B was composed of 0.1 % FA and 5% DMSO in acetonitrile. Peptides were eluted into the mass spectrometer using an electrospray ionization source (DuoSpray™ Source, ABSciex^®^) with a 25 µm internal diameter hybrid PEEKsil/stainless steel emitter (ABSciex^®^). The ionization source was operated in the positive mode set to an ion spray voltage of 5 500 V, 25 psi for nebulizer gas 1 (GS1) and 25 psi for the curtain gas (CUR).

For DDA experiments, the mass spectrometer was set to scan full spectra (m/z 350-1250) for 250 ms, followed by up to 100 MS/MS scans (m/z 100–1500) per cycle to maintain a cycle time of 3.309 s. The accumulation time of each MS/MS scan was adjusted in accordance with the precursor intensity (minimum of 30 ms for precursor above the intensity threshold of 1000). Candidate ions with a charge state between +2 and +5 and counts above a minimum threshold of 10 counts per second were isolated for fragmentation and one MS/MS spectrum was collected before adding those ions to the exclusion list for 25 seconds (mass spectrometer was operated by Analyst^®^ TF 1.7, ABSciex^®^). The rolling collision energy (CE) was used with a collision energy spread (CES) of 5.

For SWATH-MS-based experiments, the mass spectrometer was operated in a looped product ion mode [16] and the same chromatographic conditions were used as in the DDA experiments described above. A set of 60 windows of variable width (containing 1 m/z for the window overlap) was constructed, covering the precursor mass range of m/z 350-1250. A 250 ms survey scan (m/z 350-1500 m/z) was acquired at the beginning of each cycle for instrument calibration and SWATH-MS/MS spectra were collected from the precursors ranging from m/z 350 to 1250 for m/z 100–1500 for 20 ms resulting in a cycle time of 3.304 s. The CE for each window was determined according to the calculation for a charge +2 ion centered upon the window with variable CES according to the window.

### Mass spectrometry data processing

A specific library of precursor masses and fragment ions was created by combining all files from the DDA experiments and used for subsequent SWATH processing. Libraries were obtained using ProteinPilot™ software (v5.1, ABSciex^®^), using the following parameters: i) search against a database from SwissProt composed by Homo Sapiens (released in March 2019), and MBP-GFP [15] and horse ferritin light and heavy chains sequences ii) acrylamide or iodoacetamide alkylated cysteines, for whole serum or HMW respectively, as fixed modification; iii) trypsin as digestion enzyme and iv) urea denaturation as a special factor in the case of the HMW samples. An independent False Discovery Rate (FDR) analysis using the target-decoy approach provided with Protein Pilot software was used to assess the quality of the identifications, and positive identifications were considered when identified proteins and peptides reached a 5% local FDR [17, 18].

Data processing was performed using the SWATH™ processing plug-in for PeakView™ (v2.0.01, AB Sciex^®^) [19]. After adjustment of retention time using a combination of IS and endogenous peptides, up to 15 peptides, with up to 5 fragments each, were chosen per protein, and quantitation was attempted for all proteins in the library file identified from ProteinPilot™ searches.

Protein levels were estimated based on peptides that met the 1% FDR threshold with at least 3 transitions in at least six samples in a group, and the peak areas of the target fragment ions of those peptides were extracted across the experiment using an extracted-ion chromatogram (XIC) window of 5 minutes with 100 ppm XIC width. Protein levels were estimated by summing all the transitions from all the peptides for a given protein (an adaptation of [20] and further normalized to the levels of the IS [15]).

### Statistical analysis and biological interpretation

Pearson’s Chi-squared Test for Count Data was performed in R version 4.2.1, using the chisq.test function available in the native stats package in R to determine if there were significant differences in the gender proportion of the groups within the studied cohort.

To assess the variation of the serum proteins (either Whole serum or the HMW fraction of the serum) among the three groups, a Kruskal–Wallis H test was followed by the Dunn’s Test for pairwise comparison. Dunn’s p-values were corrected using the Benjamini-Hochberg FDR adjustment, and statistical significance was considered for p-values below 0.05.

Stepwise Linear Discriminant Analysis (LDA) was performed to select the proteins responsible for the best separation of the groups being studied. LDA was performed using IBM^®^ SPSS^®^ Statistics Version 22 (Trial). LDA was attempted considering the proteins only altered at the Whole Serum or HMW Serum, and for the combination of both results. The evaluation of the models obtained from each analysis was performed by comparison of the Receiver operating characteristic (ROC) curves obtained using each model. A comparison of the ROC curves was performed using MedCalc Statistical Software version 20.106 (MedCalc Software Ltd;[21]; Trial). The Delong et al. (1988) [22] method was used for the calculation of the Standard Error (SE) of the Area Under the Curve (AUC) and of the difference between two AUCs, and the Confidence Interval (CI) for the AUCs were calculated using the exact Binomial Confidence Intervals which are calculated as the following AUC ± 1.96 SE.

Violin plots were used to present the distribution of the individual protein levels among each condition, and Pearson’s correlation analysis was performed to evaluate the similarity between the profiles of the proteins highlighted in the study. Violin plots were generated using GraphPad Prism 8.0.1 (Trial) and the Pearson’s correlation was performed using Morpheus software [23]. Heatmap and hierarchical clustering analyses were computed using PermutMatrix version 1.9.3 [24, 25] using the Euclidean distance and McQuitty’s criteria.

Physical protein-protein interactions between the highlighted analytes were predicted by GeneMANIA webserver (Gene Function Prediction using a Multiple Association Network Integration Algorithm; [26, 27] together with a gene ontology (GO) analysis of the formed network. In addition to the proteins imported from this study, 28 additional related genes were allowed to create the interaction network using equal weighting by network. An additional GO enrichment analysis considering the term “biological process” was also performed. On the other hand, functional protein association networks were evaluated using the Search Tool for Retrieval of Interacting Genes/Proteins (STRING) version 11.5 [28] with a medium confidence of 0.4 [29].

Pathway enrichment analyses were performed using the FunRich software (version 3.1.3) [30], considering two different databases: the FunRich or the Reactome database. In both instances, a statistical analysis employing a hypergeometric test was conducted, using the FunRich human genome database as the background. Enriched pathways were considered for a non-corrected or a Bonferroni-corrected p-valuelllbelowlll0.05 for Funrich or Reactome database, respectively.

## Results

### High molecular weight fractionation as a reproducible method for biomarker discovery

To explore the potential of HMW fractionation in peripheral biomarker research, serum samples underwent either ultrafiltration using a 300 kDa MWCO filter followed by protein precipitation of the retentate (referred to as HMW serum), before protein digestion and MS analysis, or were directly analyzed (whole serum) (see Figure 1). In the proposed pipeline, the HMW fractionation was performed under non-denaturing conditions, such that the filter would retain native HMW protein complexes and large molecular structures.

**Figure 1.**
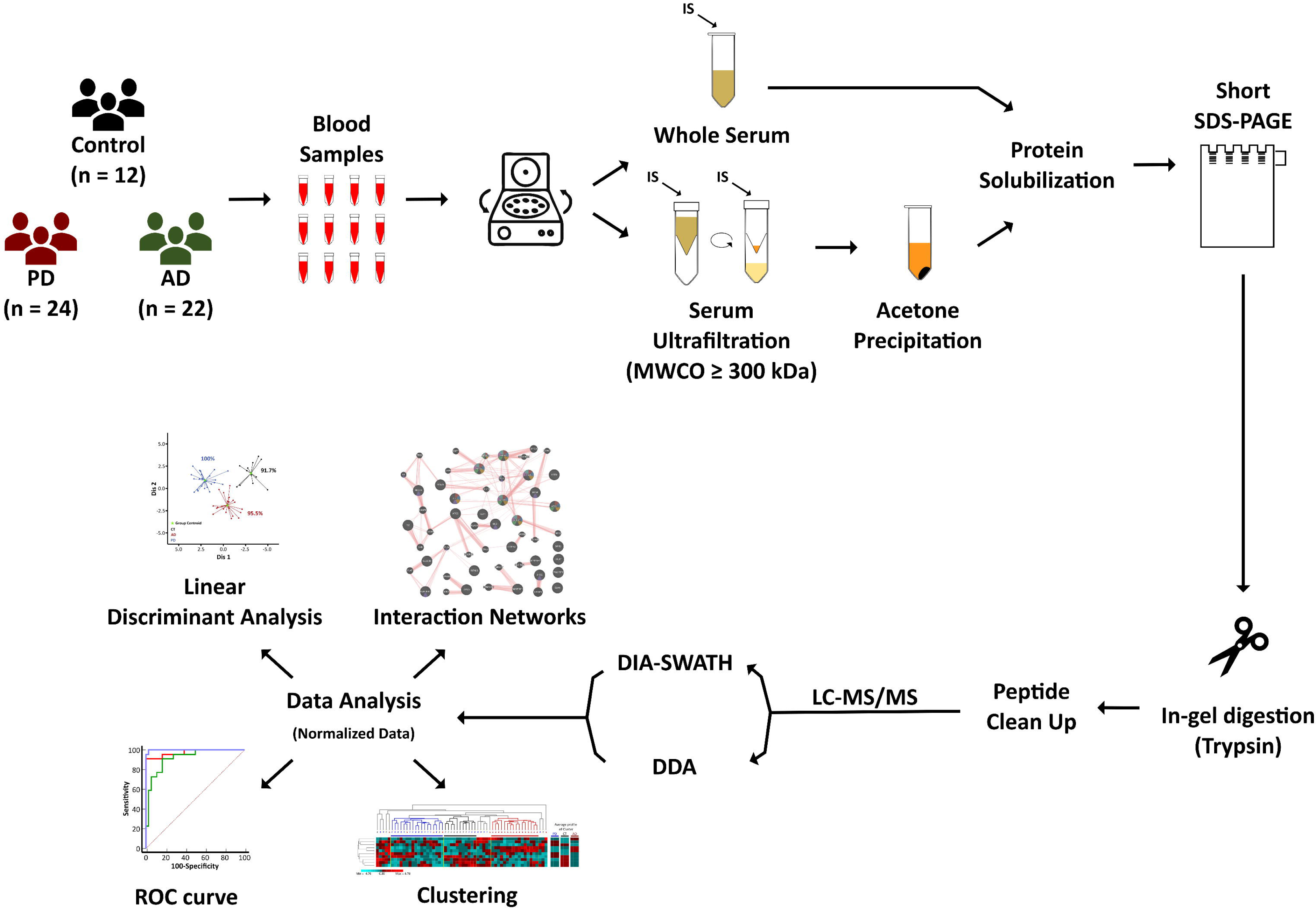
Pipeline of sample preparation, data acquisition and data analysis.

As previously reported, employing an appropriate IS is crucial for delineating effective proteome changes among different groups [12]. Given the nature of the fractionation proposed in this study, an ideal IS for this procedure would be a HMW protein that remains retained by the filter and bears no similarity to other human proteins. In this context, equine globular protein Ferritin (∼440 kDa size) emerged as a suitable IS for several reasons: it lacks similarity with any other protein from the human proteome (Additional File 1: Supplementary Figure 1a), the peptides monitored in the SWATH-MS analysis are easily distinguishable from the matrix (the human serum proteome; Additional File 1: Supplementary Figure 1b), and it exhibits a coefficient of variation similar to the one obtained by the MBP-GFP used in the unfractionated analysis and previously characterized [12] (Additional File 1: Supplementary Figure 1c). Additionally, the overall reproducibility of the fractionation was also inspected. Similarly, to what was observed for the IS (Additional File 1: Supplementary Figure 1c), the overall coefficient of variation of the proteins quantified using technical replicates (Additional File 1: Supplementary Figure 1d) revealed that this procedure did not induce an appreciable increase in the variability of the quantification when compared with the conventional protocol (unfractionated samples). Moreover, the variation caused by the sample processing steps may be reverted by the normalization of the values to the IS since the coefficient of variation of the IS is similar to the one observed for the majority of the proteins, indicating that the selected IS is a good predictor of the alterations induced by the method.

### High molecular weight fractionation increases the detection of proteomic alterations in NDs

As proof of concept, this procedure was applied to serum samples of a cohort comprised of 3 different groups: AD (n = 22), PD (n = 24), and CT (n = 12). No statistically significant differences were found in the gender distribution between the three groups; however, differences were observed concerning age distribution, with PD individuals being slightly younger, on average, than both other groups (Table 1).

**Table 1.** Study population distribution across age and sex.

|  | AD<br>(n = 22) | PD<br>(n = 24) | CT<br>(n = 12) | p-value |  |  |  |  |
| --- | --- | --- | --- | --- | --- | --- | --- | --- |
|  |  |  |  | CT vs.<br>Diseases | CT vs. AD vs. PD | AD vs. PD | CT vs. PD | CT vs. AD |
| **Male (n)** | 9 | 11 | 3 | 0.197<sup>*</sup> | - | 0.643<sup>*</sup> | 0.148<sup>*</sup> | 0.262<sup>*</sup> |
| **Female (n)** | 13 | 13 | 9 |  |  |  |  |  |
| **Age**<br>**(mean ±**<br>**SD)** | 68.4 ±<br>8.2 | 60.3 ±<br>10.4 | 67.5 ±<br>7.2 | - | 0.014<sup>§</sup> | 0.009<sup>#</sup> | 0.033<sup>#</sup> | 0.396<sup>#</sup> |

<sup>\*</sup> Determined by $\chi^2$ test according to each group's male/female proportions. <sup>§</sup> Determined by Kruskal-Wallis rank sum test to compare age between all groups. <sup>#</sup> Determined by Dunn's test (Benjamini-Hochberg correction) to compare age between group pairs. Diseases refer to AD and PD grouped together.

Patients with neurodegenerative diseases, both AD and PD, were selected for this study. These conditions are frequently associated with the development of abnormal protein complexes and protein aggregation. Hence, they offer a fitting context to evaluate the fractionation approach’s effectiveness in exploring potentially altered protein interactions—an aspect frequently neglected in conventional proteomics analyses.

A total of 203 and 186 proteins were quantified in whole serum and HMW serum, respectively (Figure 2a, solid lines; Additional File 2: Supplementary Tables 1 and 2 for detailed information). A large overlap was observed between both sample preparation procedures (168 proteins were shared, which corresponds to more than 70% of all the quantified proteins). Additionally, quantification in the whole serum of the proteins identified in the HMW serum library did not lead to a discernible increase in proteome coverage (Figure 2a, dashed line; Additional File 2: Supplementary Table 3). Altogether, these results indicate that there are no major differences in terms of the proteins being quantified in the two approaches, revealing that the main aim of the HMW fractionation presented in this work – fractionation under non-denaturing condition – is not the overall improvement of the proteome coverage but the possibility of interrogating the samples considering protein interactions/macromolecular organization.

**Figure 2.**
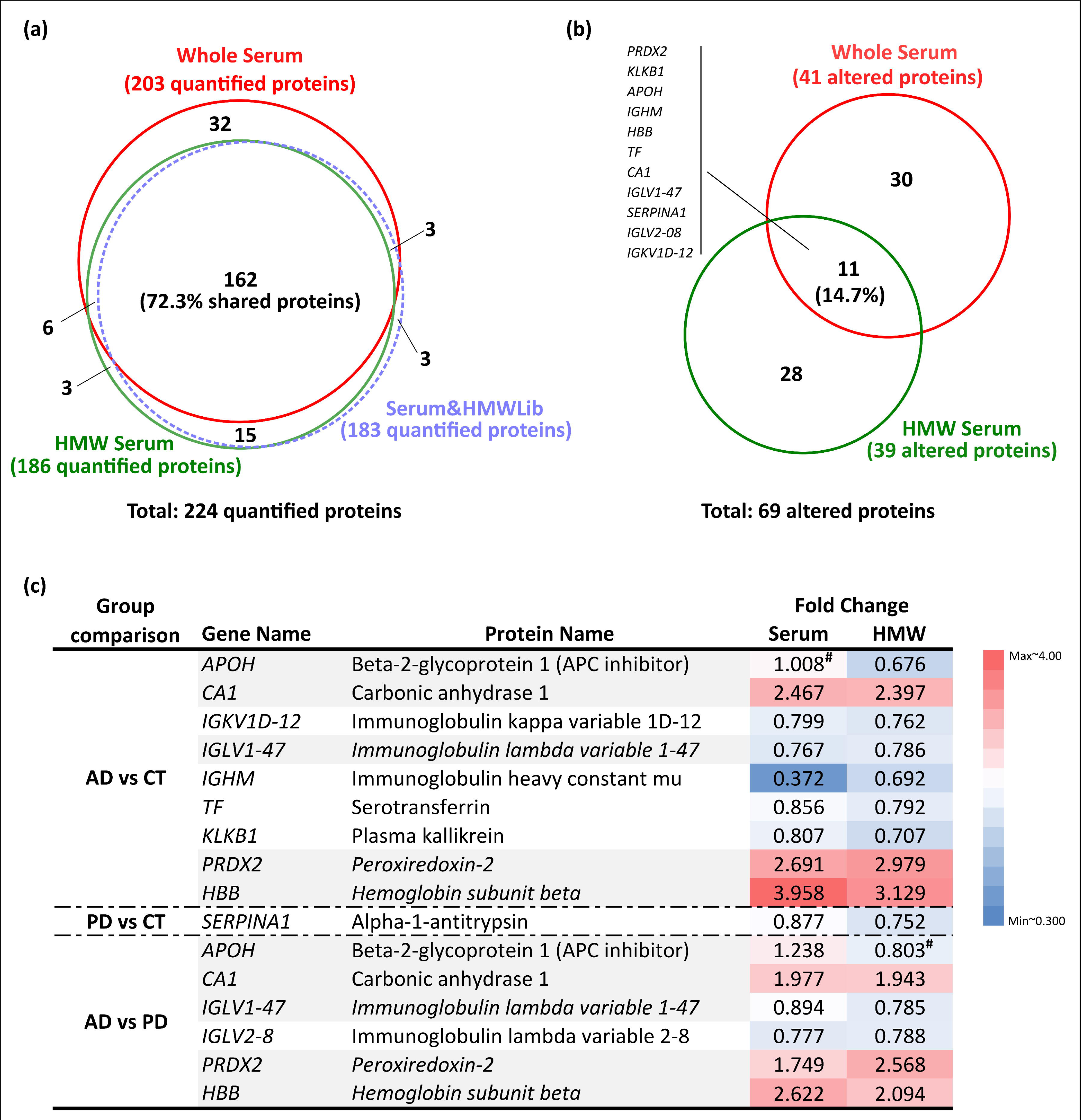
Comparative differential proteomic analysis of whole and HMW serum of AD and PD patients. **(a)** Venn diagram comparing the number of quantified proteins in each sample type using sample-specific libraries (solid lines; Additional File 2: Supplementary Tables 1-2). In addition, quantification in whole serum was also performed for the proteins identified in the HMW-specific library (dashed line; Serum&HMWLib condition; Additional File 2: Supplementary Table 3) to evaluate the possible use of HMW fractionation as a tool to improve the proteome coverage of serum samples. A total of 224 proteins were quantified in the serum samples using the three strategies referred above; from those, 162 proteins were commonly quantified independently of the strategy, corresponding to nearly three-quarters of all the quantified proteins (72.3%). Only three new proteins were quantified in whole serum using the HMW-specific library. **(b)** Venn diagram comparing the total number of proteins considered altered among the three experimental groups using the two different serum-processing strategies used in this work (Additional File 1: Supplementary Figure 2 and Additional File 2: Supplementary Tables 1-2). A total of 69 proteins were considered altered among the three experimental groups; of those, only 11 proteins (the respective gene names are indicated) were consistently considered altered independently of the strategy used. **(c)** Comparison of the levels of the 11 proteins commonly considered altered in both whole serum and HMW-fractionated serum. The proteins were arranged considering the group comparison where the statistical differences were observed (Additional File 1: Supplementary Figure 3), and the alterations were presented as the median fold change observed in each sample type. Proteins considered altered in more than one group are indicated in italic and with a grey shadow. Only one protein, the beta-2-glycoprotein 1 (indicated in bold), presented a divergent tendency when considering its values in the whole serum versus after the HMW fractionation of the samples. # - non-statistically significant difference.

This was further confirmed by the fact that only 14.7% of all the proteins considered altered in this study (11 out of 69 proteins) were consistently altered in the fractionated and unfractionated samples (Figure 2b; Additional File 2: Supplementary Tables 1-2). Moreover, taking into consideration the changes of those 11 commonly altered proteins, it is possible to observe that all proteins, except for the protein beta-2-glycoprotein 1 (Figure 2c, bold), presented the same tendency in both fractionated and unfractionated samples (Figure 2c). These results, in combination with the previous observations, demonstrate that, although the two approaches quantified the same set of proteins, each method interrogated the samples in a particular context, resulting in the identification of a different set of altered proteins and the possibility to identify different regulatory mechanisms: while the direct analysis of the serum mainly represents the alteration at the protein level, the HMW fractionation under non-denaturing conditions may allow the evaluation of the physical interaction of the proteins.

These results are further supported by the analysis of the MW distribution of the proteins being studied, which reveals a similar profile between both approaches (Additional File 1: Supplementary Figure 4a, 4b, and 4c), with most of the proteins detected in HMW serum presenting a MW below 150 kDa. Moreover, a similar distribution is observed for the proteins altered exclusively in the HMW approach (Additional File 1: Supplementary Figure 4d), indicating that this strategy is not biased towards only the HMW proteins and supporting the idea that those proteins altered in this fraction may correspond to proteins being organized into different complexes.

### High molecular weight fractionation revealed differences in the macromolecular organization of the serum proteome

To test the hypothesis that the HMW approach is capable of evaluating the re-organization of protein complexes, the nature of the 28 proteins altered exclusively in HMW serum was evaluated. This analysis revealed that most of those proteins had been described to interact physically either with each other or with other proteins (Figure 3a). Upon immediate observation of the GeneMania network, which primarily evaluates the interactions between the proteins under study (Additional File 2: Supplementary Table 4), it becomes apparent that the majority of the proteins participate in established physical interactions. Moreover, it can be observed that these proteins form a large and interconnected network comprising 20 of the 28 altered proteins centered around the interactions between the apolipoproteins and lecithin-cholesterol acyltransferase (encoded by the LCAT gene). Although not participating in any known interactions with another protein from the 28 altered proteins, the proteins Glutathione peroxidase 3 (GPX3), Pigment epithelium-derived factor (SERPINF1), Thyroxine-binding globulin (SERPINA7), and Alpha-1B-glycoprotein (A1BG) are also known to establish physical interactions including with the proteins identified in this analysis. Only four of the 28 altered proteins, namely SRR1-like protein (SRRD), carnosine dipeptidase 1 (CNDP1), peptidoglycan recognition protein 2 (PGLYRP2), and serum amyloid A-4 protein (SAA4), were found to have no disclosed interactions in this particular analysis. However, interactors for those proteins have already been pointed out in some screening assays, as confirmed in BioGRID (Biological General Repository for Interaction Datasets, Additional File 2: Supplementary Table 5). Furthermore, as revealed by the functional analysis, several of these 28 proteins are involved in the formation of complexes with lipids and platelet components (Additional File 2: Supplementary Table 6), indicating that those proteins can form complexes not only via the interaction with other proteins but also with other molecules, and thus be organized in large complexes. This involvement in the potential formation of macromolecular complexes is even more evident when the functional pathways enriched in each of the two lists of proteins (30 and 28 altered proteins exclusively in the whole serum or HMW serum, respectively) are directly compared (Figure 3b). This comparison highlights the fact that all the pathways that are either only, or at least more, enriched in the HMW dataset in comparison to the whole serum dataset (pathways indicated in bold) are related to the formation/regulation of large complexes/macrostructures, namely amyloids, fibrin clot formation and dissolution pathways and lipoprotein-related metabolism. On the other hand, the pathways that are particularly enriched or unique in the whole serum dataset are mainly related to transcription factor networks (Additional File 2: Supplementary Table 7).

**Figure 3.**
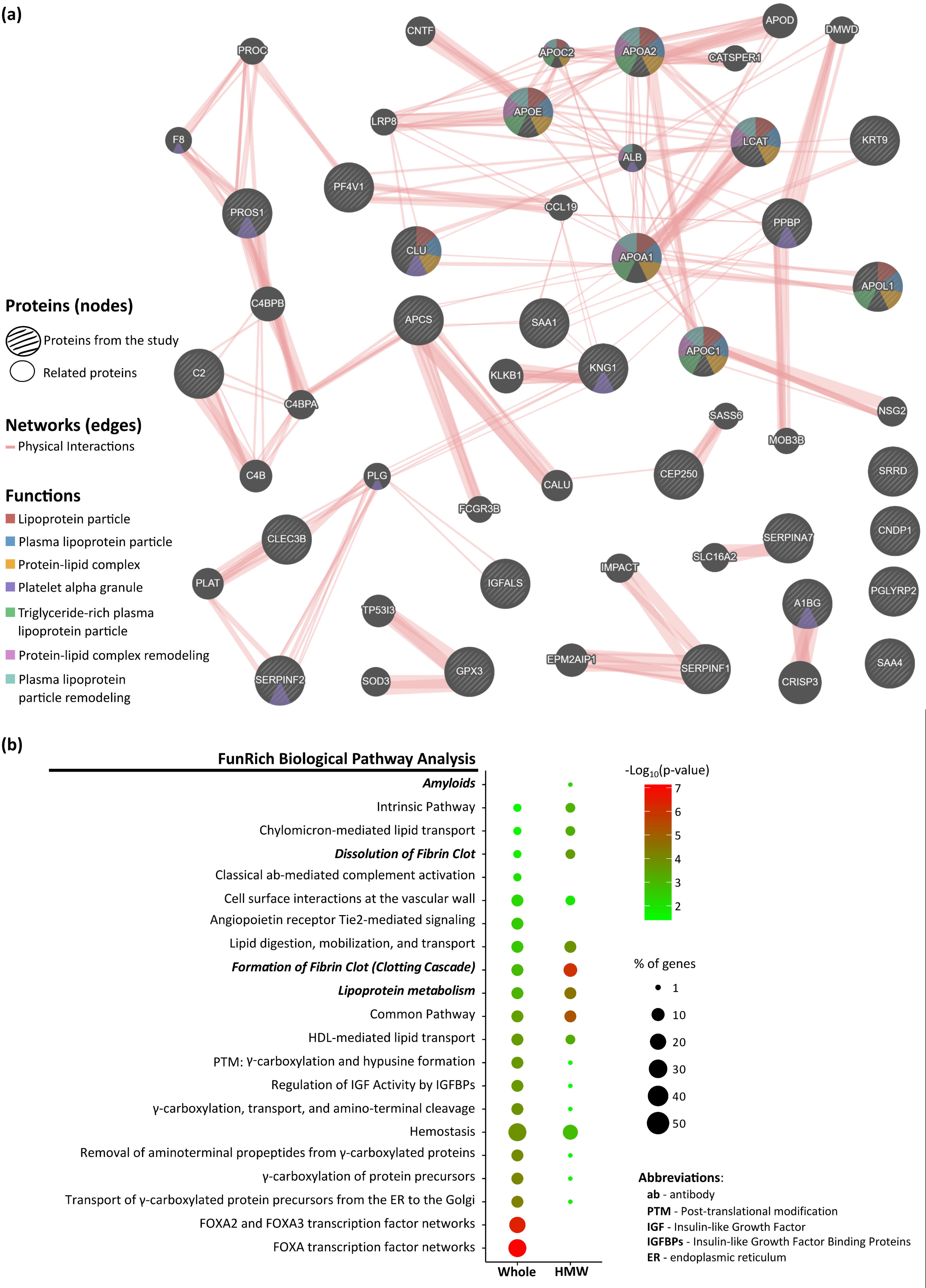
Characterization of the proteins exclusively altered in whole serum or HMW serum. **(a)** GeneMania Network of the 28 proteins altered only at the HMW serum sample (listed in larger circles). The analysis was performed with network weighting equal by network, allowing a maximum of 28 extra resultant genes (non-listed small circles). The seven most enriched GeneMania Functions were highlighted in the network (color code; complete results in Additional File 2: Supplementary Table 4). Only protein-protein physical interactions (red edges) were considered in this analysis, demonstrating that most of these 28 altered proteins have known interactors and can be involved in the formation of large protein-protein complexes. **(b)** FunRich Biological Pathways enriched in the whole serum (30 proteins; Additional File 2: Supplementary Table 5) and HMW serum (28 proteins; Additional File 2: Supplementary Table 6) proteomes. All GO analyses considered a p <0.05. Pathways uniquely enriched at the HMW serum or particularly enriched in this type of sample when compared to the whole serum are indicated in bold.

### Evaluation of the potential of this combined strategy for biomarker discovery

The previous set of results demonstrates that both approaches can provide complementary information. In line with this evidence, the potential to use this combined strategy for biomarker discovery was also evaluated. In general, both approaches result in nearly 40 proteins altered in at least one pair of comparisons (Figure 4a-b; Additional File 1: Supplementary Figure 3 for details regarding each pair of comparisons), with a tendency to have more proteins altered in the comparisons involving the AD group (at least 20 proteins were altered compared to CT against a maximum of 14 altered proteins in PD vs. CT, Additional File 1: Supplementary Figure 2a-b) and only a small subset of proteins altered in only one comparison (15/41 in the whole serum and 11/39 in HMW serum, Figure 4a and 4b, respectively). Besides those similarities, different profiles of altered proteins were observed depending on the approach used. Hence, it can also be highlighted that while the whole serum strategy (Figure 4a) mainly found proteins altered between the two disease groups (39 proteins in AD vs. PD compared to 20 and 12 proteins in AD vs. CT and PD vs. CT, respectively), the HMW approach (Figure 4b) captured more differences between AD and CT samples (36 out of the 39 altered proteins). Due to this complementarity, the combination of results from both approaches resulted in a more comprehensive profile (Additional File 1: Supplementary Figure 2a-c), with a general increase in the number of altered proteins per group (total of 69 altered proteins, Figure 4c). This improvement is particularly evident in the case of proteins altered between PD and CT samples, for which only one protein was considered commonly altered in the two approaches (Figure 2c and Additional File 1: Supplementary Figure 2b), thus resulting in the duplication of the list of proteins with the potential to serve as biomarkers for PD versus CT individuals. Common to all mapped profiles was the low number of proteins altered between all three groups (4, 1, and 6 for Whole serum, HMW serum and the combination of both, Figure 4a-c, respectively) and the absence of proteins altered exclusively between PD and CT samples.

**Figure 4.**
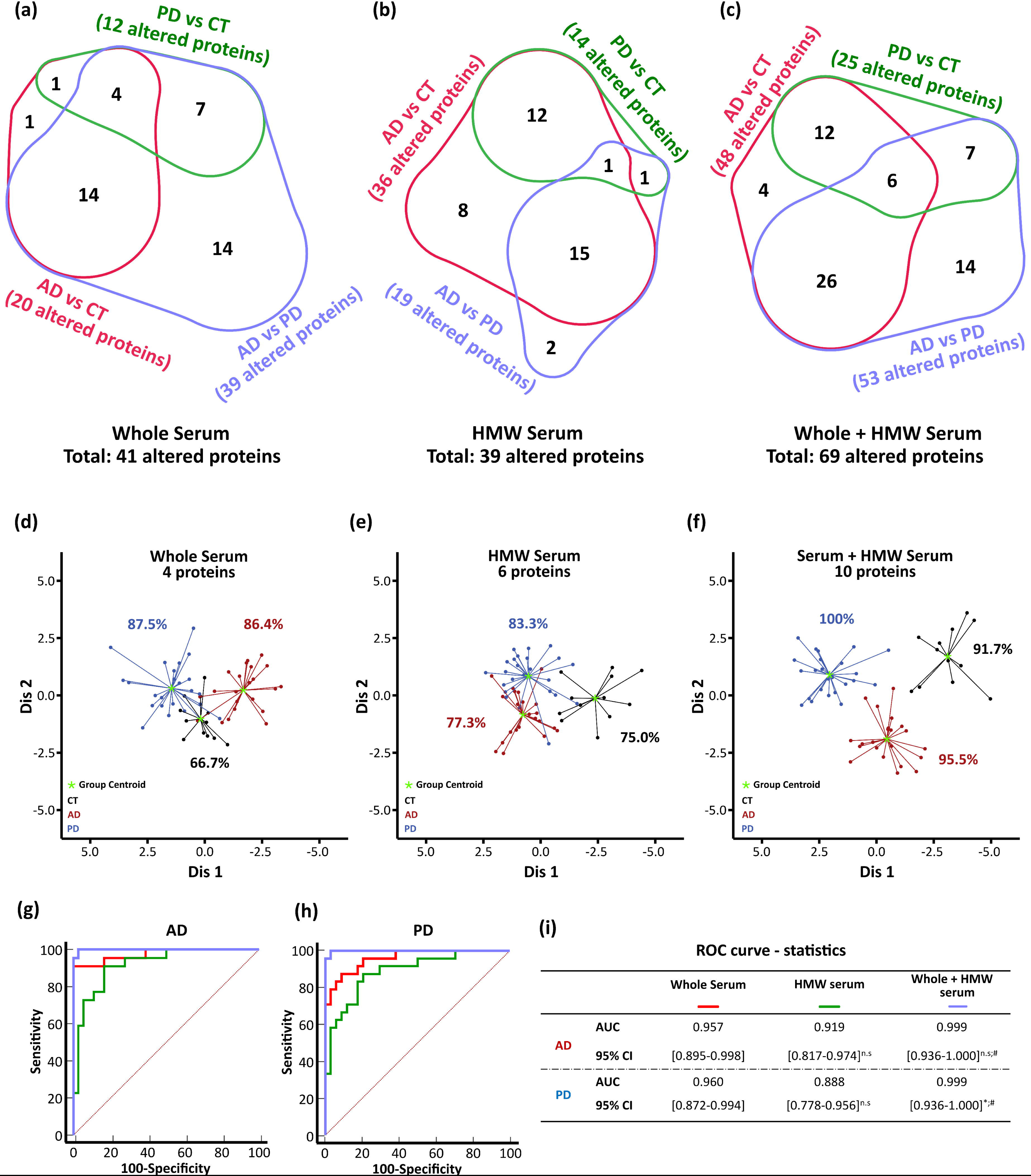
Identification of potentialcirculating biomarkers of AD and PD. **(a - c)** Venn diagrams representing the distribution of the altered proteins among the different comparisons (AD vs. CT; PD vs. CT; AD vs. PD) considering the whole serum, HMW serum and the combination of the two (Whole + HMW serum) types of samples, respectively. **(d-f)** LDA using all altered proteins per sample type or the combination of the two. The number of proteins used in each model is indicated above the graphic, and for each model it is indicated their specificity (percentage of healthy individuals correctly identified; indicated in black) and the sensitivity per disease condition (percentage of AD or PD patients correctly identified; indicated in red and blue, respectively). Specificity and sensitivity values were also summarized in Additional File 2: Supplementary Table 7, and the generated LDA discriminant functions and their respective statistical confidence were summarized in Table 2. **(g-h)** Comparative ROC curves of the discriminant functions generated using all altered proteins per sample type or a combination of the two. An independent evaluation was performed for each disease group being studied. The AUC of the ROC curves, their 95% CI and the pairwise comparisons are summarized in **(i)**. CI was calculated as follows AUC ± 1.96 SE. n.s., non-significant alterations. * and # indicate a plll<lll0.05 for statistically significant differences in comparison to the Whole Serum or HMW+Whole Serum in comparison to HMW Serum, respectively, using the method of Delong et al. (1988) [22].

To further confirm the biomarker potential of the altered proteins from the three strategies presented above, they were used as input to build discriminant models that could differentiate between the studied groups (Figure 4d-f). From each dataset, candidates whose combination resulted in the best possible discriminant model were automatically selected (detailed information regarding the methods in Table 2), resulting in three distinct and statistically valid models (all with p < 0.0032) capable of discriminating the three groups being studied. The whole serum dataset resulted in a reasonable model composed of 4 proteins (Figure 4d), with a median sensitivity (the capacity to classify the individuals from each disease group correctly) of 86.95% (sensitivity and specificity are summarized in Additional File 1: Supplementary Table 8). On the other hand, the model created with six proteins from the HMW approach (Figure 4e) had lower performance, with a median sensitivity of only 80.3% (corresponding to 83.3% predicting capacity for PD samples and 77.3% for AD samples). Moreover, neither of the models was particularly good in the classification of CT samples, resulting in a specificity of only 66.7% and 75% in the whole serum and HMW serum models, respectively. Remarkably, the combined model (created from the dataset containing the altered proteins from both approaches - Figure 4f) clearly outperformed the two models based only on proteins from a single approach. For this combined model, 10 proteins were selected and integrated, creating a discriminant model capable of correctly classifying more than 96% of all tested samples (93% in a cross-validation test; Table 2). This included the correct classification of 100% of samples from the PD group (Figure 4f), which encompassed samples from younger patients compared to the other studied groups (Table 1). However, it was confirmed that this age difference did not interfere with the classification of samples when employing the discriminant model since: i) age by itself did not yield good classification performance (Additional File 1: Supplementary Figure 5) and ii) none of the 10 proteins included in the combined model were strongly correlated with age (Additional File 1: Supplementary Figure 6). Altogether, this combined method presented a median sensitivity of 97.75% and a specificity of 91.7%.

**Table 2.**
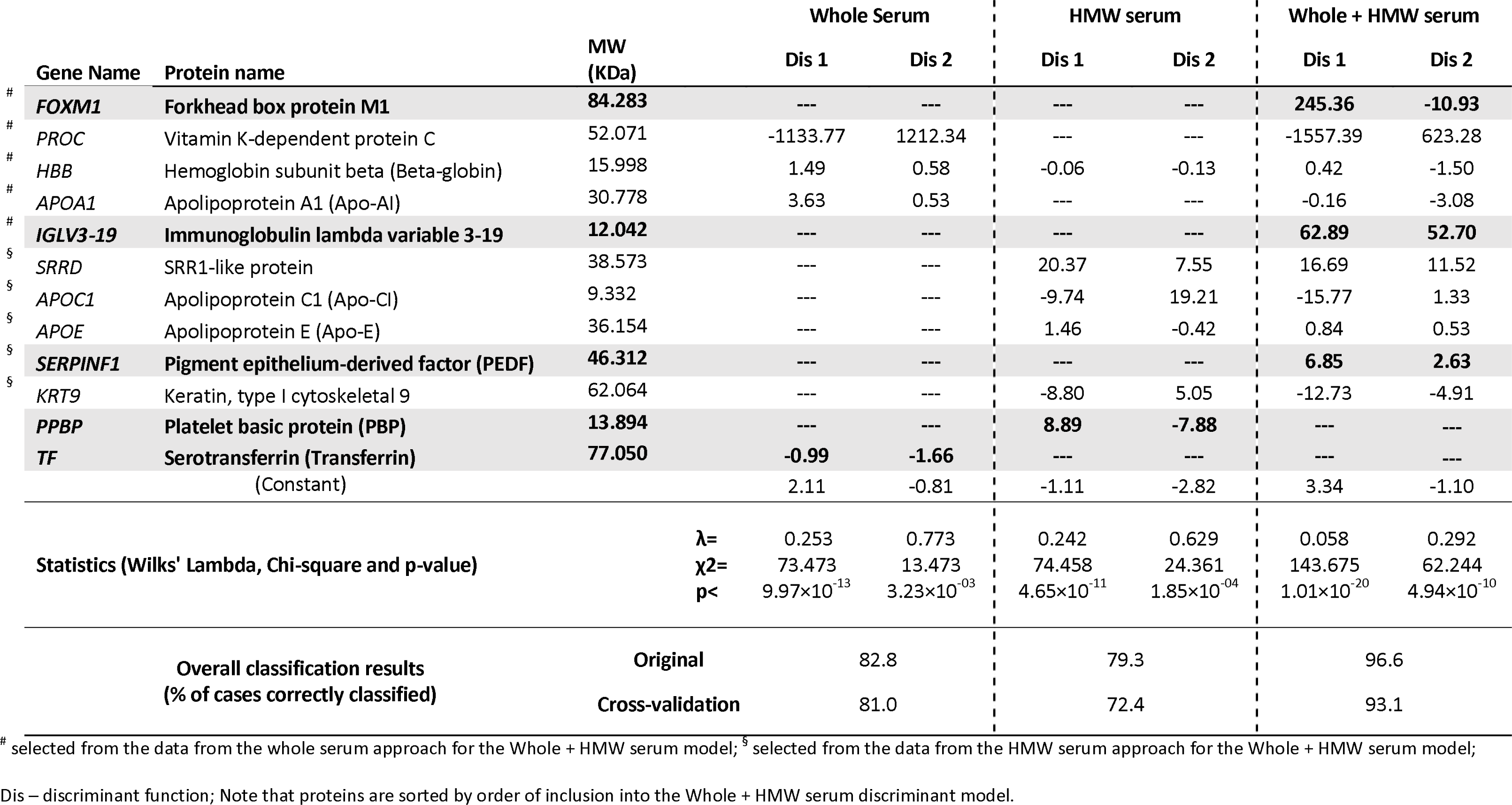
Linear Discriminant models, respective statistical analysis and classification results.

The diagnostic capacity of these models was further assessed by ROC curves, which measured their ability to accurately classify AD and PD patients from among all other samples (Figure 4g-i). This analysis confirmed that the best model is the one created with the combination of proteins from both approaches and that, in general, the model using only proteins from the whole serum approach is better than the model from the HMW approach. The respective statistics (Figure 4i) further support that the whole serum model performed better than the HMW serum model but without statistically significant differences between the two ROC curves. Additionally, the statistical analysis also confirmed that the combined model (AUC = 0.999 for the classification of AD and PD patients) is the best model and that it performed significantly better (p < 0.05) than both other models for PD classification (HMW serum, AUC = 0.888; whole serum, AUC = 0.960) and better than the HMW serum model (AUC = 0.919) in the case of AD classification. The robustness of the combined model is further evidenced by the confidence interval (CI) of the AUC, which has a lower limit above 0.93 for both diseases, in contrast with the values achieved for the other two methods, whose lower limits are all below 0.9.

By looking at the proteins selected to build the different methods (Table 2), it was observed that the combined method is not the simple combination of the proteins previously selected from each of the individual methods. The combined model is built by the combination of 10 proteins, five from each dataset, including three [forkhead box protein M1 (*FOXM1*), immunoglobulin lambda variable 3-19 (*IGLV3-19*) and pigment epithelium-derived factor (*SERPINF1*)] that were not selected on the database-specific models. On the other hand, some previously selected proteins [serotransferrin (*TF*) and platelet basic protein (*PPBP*)] were not included in the combined model. Finally, the Hemoglobin subunit beta (*HBB*) was selected in both approach-specific models, although only data from the whole serum dataset was used in the combined model. These results demonstrate that the increase in the initial amount of data provided for the discriminant analysis has an important impact on the generated model by making it possible to test different combinations of proteins and, thus, allowing for the identification of better combinations than those highlighted in the analysis of individual datasets. Interestingly, all ten proteins selected in the combined model have a MW below 90 kDa (Table 2). This finding confirms that all the proteins selected from the HMW approach have a MW below the theoretical cut-off of the filters used for fractionation. This observation supports the hypothesis that this approach might effectively assess the remodeling of molecular complexes. By plotting the individual values of each of the ten proteins selected in the combined model (Figure 5a), it is possible to observe that, as expected, those values present some variation characteristic of the individuality of each patient. Nevertheless, considering that the model was able to correctly classify more than 90% of all the patients (Figure 4f), it is possible to infer that the combinations performed in the model could diminish the impact of the biological variability, proving that the combination of different markers can overcome their individual weaknesses. The analysis of these results immediately reveals that i) only three proteins from the model (the proteins encoded by the genes *FOXM1*, *HBB,* and *SRRD*) are significantly altered between all three groups and ii) only one protein, the apolipoprotein C1 (Apo-CI, encoded by the *APOC1* gene), is altered between a single comparison, in this case between AD and PD which may indicate that this protein may have a particularly important role in this model for distinguishing AD from PD patients. Among the remaining 6 proteins: i) three, apolipoprotein E, pigment epithelium-derived factor and KRT9, are altered between both disease groups and control sample (all three found in the HMW fraction); ii) two, apolipoprotein A1 and immunoglobulin lambda variable 3-19, are altered in AD in comparison to both PD and CT, and one, the Vitamin K-dependent protein C (encoded by the PROC gene), is altered in PD patients in comparison to the other two groups. Moreover, it is noteworthy that while there was a tendency to incorporate proteins that were increased in AD compared to CT from the whole serum approach (three out of the five proteins from the whole serum model, encoded by the genes *FOXM1*, *HBB*, and *APOA1*), the opposite trend was observed in the case of proteins from the HMW approach. Specifically, three out of the five proteins (the proteins encoded by the genes *SRRD*, *APOE*, and *SERPINF1* ) were found to be decreased in the AD vs. CT comparison. On the contrary, for the PD vs. CT comparison, there were no major differences in terms of tendencies when considering the proteins captured in the whole serum or the HMW serum.

**Figure 5.**
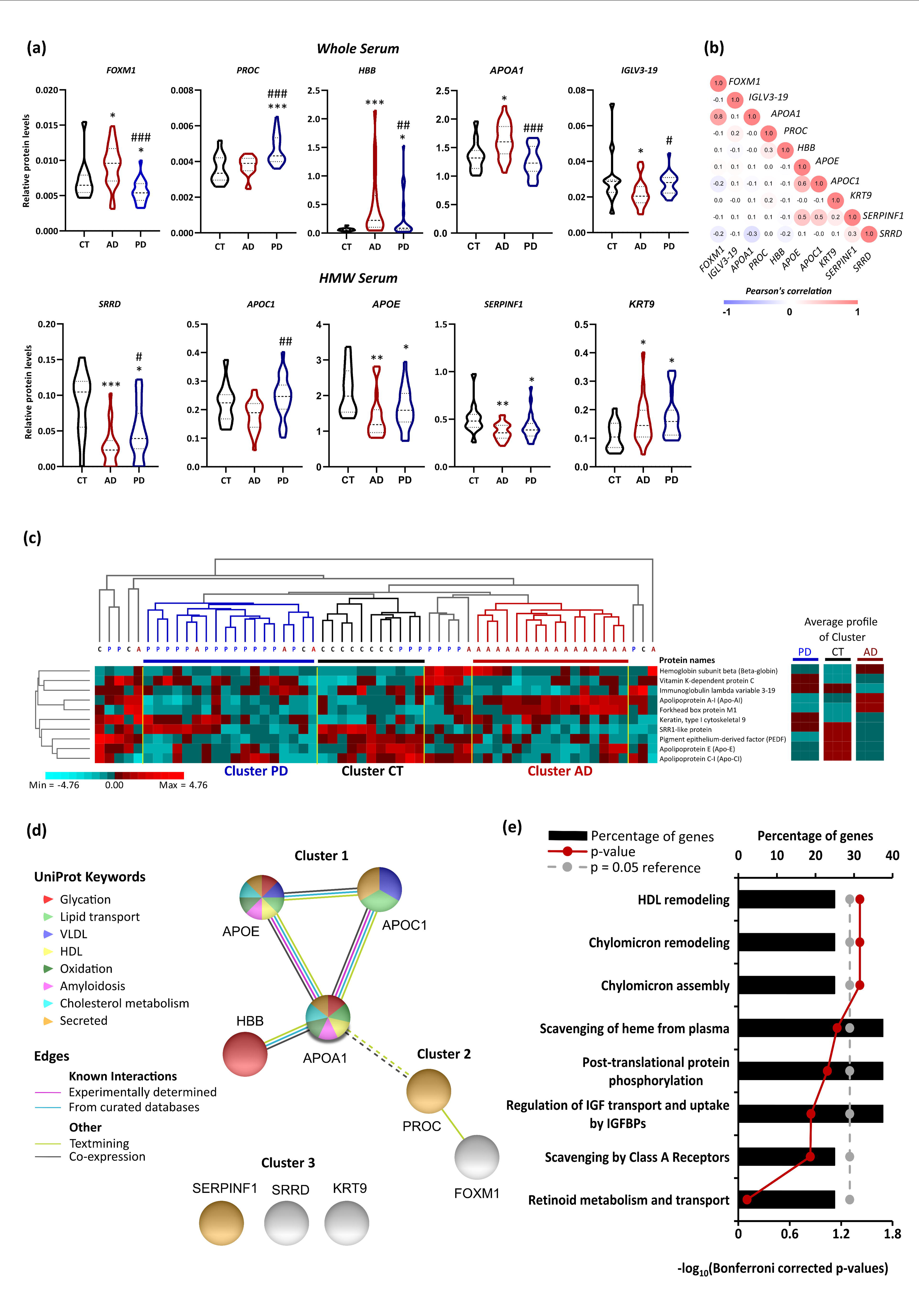
Characterization of the protein panel selected by the combined LDA model. **(a)** Violin plots representing the protein level distribution of the ten proteins from the model. The dashed lines indicate the first, second (median) and third quartiles. *, **, and *** indicate a plll<lll0.05, p<0.01, and p<0.001 for statistically significant differences. * denotes the comparison to control and # the differences between disease groups. Statistical analysis was performed using the Kruskal–Wallis H test followed by the Dunn’s Test for pairwise comparison. **(b)** Pearson’s correlation analysis between the overall regulation profile of the proteins included in the model. **(c)** Heatmap and hierarchical clustering analysis of the proteins from the model. Clustering was performed for both proteins and individuals. Three different clusters (Cluster PD, CT and AD) containing the large majority of the individuals from a given group can be highlighted from the analysis. The average profile of each cluster is indicated on the right and can be considered as the profile of expression of those proteins within the groups considered in this study. **(d)** Interaction network of these proteins carried out with STRING with a medium confidence (0.4) score. The color of the edges indicates the type of evidence that supports the interaction, while the color of the nodes represents the categorization of the proteins considering UniProt Keywords. The calculated PPI enrichment p-value is 2.07e-05. Three clusters (Cluster 1 to 3) can be identified within the network, with the dashed edges indicating the separation between clusters. Cluster 1 corresponds to proteins whose interactions are experimentally confirmed, cluster 2 is composed of theoretically related proteins, and cluster 3 corresponds to non-related proteins. **(e)** Reactome pathways enrichment analysis using the proteins from the model. The red line indicates Bonferroni’s corrected p-value with the corrected *p* < 0.05, meaning a significant enrichment. The grey dashed line indicates the reference line (*p* = 0.05).

Finally, all proteins from the whole serum dataset, in addition to SRR1-like protein (encoded by the *SRRD* gene) and Apo-CI from the HMW dataset, were significantly altered between both disease groups. From these, three proteins [the protein Forkhead box protein M1 (encoded by the *FOXM1* gene), hemoglobin subunit beta (encoded by *HBB* gene), and apolipoprotein A1 (Apo-AI, encoded by the *APOA1* gene)] are less abundant in PD samples than in AD samples, while the remaining four are increased.

The correlation analysis of the protein abundances among groups confirmed that, in general, there is no particularly evident correlation between the profiles and the magnitude of regulation of these proteins (Figure 5b). There were, however, some observed exceptions, including a strong positive correlation between the proteins encoded by the genes *APOA1* and *FOXM1* (r = 0.8) and, to a lesser extent, the proteins encoded by *APOC1*, *APOE and SERPINF1* (r = 0.5-0.6). Interestingly, the proteins showing a positive correlation originated from the same approach, the gene products of *APOA1* and *FOXM1* were both highlighted in the whole serum approach, suggesting that the overall levels of these two proteins were similarly altered. Meanwhile, the products of the *APOC1*, *APOE*, and *SERPINF1* genes were identified as altered in the HMW fractionation strategy, which may indicate that these proteins might be part of the same complex and thus regulated in a similar manner. No remarkable negative correlations were found, with the strongest correlation observed between the proteins encoded by the *APOA1* and *SRRD* genes, which indicates that none of the proteins in the model present a completely opposite regulation profile. Additionally, an unsupervised clustering analysis using these ten proteins (Figure 5c) confirmed their capacity to partially distinguish the three groups being studied, revealing that, besides the existence of individual variability, it was possible to identify three independent clusters composed exclusively or mainly of samples from one of the three groups. This analysis also demonstrates that this set of proteins is particularly efficient in isolating the AD patients from the remaining individuals from the study: the AD cluster was composed exclusively of AD patients, and only 6 out of the 22 AD patients were not included in this cluster. On the other hand, a slightly lower separation capacity was observed for both PD and CT samples. These two clusters contained few samples that did not belong to their respective groups, resulting in a higher percentage of individuals not properly grouped (10 out of 24 and 4 out of 12 samples for PD and CT, respectively). The discrepancies observed between the clustering analysis and the discriminant model results, where the latter correctly classified over 90% of the samples, can be attributed to the fact that the clustering analysis relied solely on the individual protein distribution profiles across the samples. In contrast, the discriminant analysis employed equations with different weightings for each protein, resulting in a single model that effectively reduces the intragroup variability while promoting a better separation between the analyzed groups. Despite that, the clustering analysis remains an important approach for understanding how the proteins are modulated within the samples. Thus, from the three different clusters highlighted in the analysis, it was possible to infer the median protein abundance profile of these proteins among the three groups. For instance, the gene products of *SERPINF1*, *APOE* and *APOC1* tend to be less abundant in both disease groups compared to CT samples. Furthermore, some proteins are more abundant in each disease group, namely the gene products of *PROC* and *KRT9* in PD samples and the gene products of *HBB*, *APOA1* and *FOXM1* in AD samples. Another disease-specific observation was the smaller amount of immunoglobulin lambda variable 3-19 (*IGLV3-19*) in AD samples compared to both other groups. Overall, these tendencies characterize the unique profiles determined for each disease group, which may be a precursor to a potential future biomarker panel that could be more informative than the analysis based on any single protein.

Finally, STRING analysis (Figure 5d and Additional File 2: Supplementary Table 9) revealed that these ten proteins have more interactions among themselves than what would be expected for a random set of proteins of the same size and degree of distribution, indicating that this set of proteins is, at least partially, biologically connected (PPI enrichment p = 2.07×10 ). This result may be mainly due to the strong network involving apolipoproteins and Hemoglobin subunit beta (cluster 1). Again, two out of the ten proteins selected for the discriminant method revealed to be associated with high-density lipoproteins (HDL) and chylomicron (ultra-low-density lipoproteins particles) remodeling and assembly (Figure 5e and Additional File 2: Supplementary Table 10), highlighting the potential importance of these mechanisms in the neurodegenerative process, and confirming that the proteins related with these mechanisms could be good biomarker candidates for their diagnosis.

Given the central role that apolipoproteins appear to play in this model, a discriminant analysis was performed using data from ten altered proteins involved in apolipoprotein-related mechanisms to investigate if the diagnostic model could be limited to this set of functionally related proteins (Additional File 1: Supplementary Figure 7). However, the generated model exhibited lower diagnostic capacity compared to the combined approach, with only 82.75% of the samples being correctly classified. The model demonstrated a specificity of 66.7%, and a sensitivity of 86.4% and 87.5% for AD and PD, respectively, yielding ROC curves with AUCs equal to or below 0.955. Thus, besides the importance of apolipoproteins, the results from these proteins alone are not enough to distinguish the three groups, emphasizing the importance of having diagnostic models based on several complementary candidates instead of a single or just a few candidates. Nonetheless, the identification of this robust core of functionally related proteins underscores the significance of the combined approach for identifying new potential biomarkers. For instance, while the dysregulation of Apo-C1 and Apolipoprotein E (Apo-E, encoded by the *APOE* gene) was discovered using the HMW fractionation approach, the dysregulation of Apo-AI was identified using the whole serum approach.

## Discussion

The present study presents a proof of concept of a novel two-pronged approach to biomarker discovery in complex peripheral biological fluids. More specifically, it was demonstrated that through the combination of two complementary proteomics strategies, the direct analysis of whole serum and the analysis of serum HMW fraction (above 300 kDa) in non-denaturing conditions, another level of proteome characterization of the samples could be achieved. This resulted in more robust diagnostic models and insights into disease mechanisms. In this sense, when applied to serum samples from a cohort of control individuals and individuals afflicted by neurodegenerative diseases, this strategy allowed for a strong discriminant model to be built, able to distinguish all studied groups more effectively than the models generated from a single proteomics analysis. The most noticeable findings from this model showed that several, otherwise overlooked, proteins may yet serve as potential biomarkers of disease, in this case, AD and PD, particularly when analyzed together in a model created using the two different approaches. Thus, these results confirm the importance of having a panel of potential candidates rather than a single protein biomarker. Furthermore, they also demonstrate that the biomarker discovery field will benefit from combining data from the sample obtained through different sample processing strategies. While not individually sufficient to be considered as biomarkers, the significant influence of apolipoproteins, particularly Apo-AI, Apo-CI, and Apo-E, in the aforementioned discriminant model, suggests a potential disease-specific dysregulation of lipoprotein metabolism in AD and PD patients.

### High molecular weight fractionation may reveal a potentially altered macromolecular and macromolecular-complex organization

In this work, it was demonstrated that interrogating serum samples with the HMW fractionation method adds an extra layer of information capable of bringing new insight into the behavior of the serum proteins, particularly regarding their potential macromolecular organization. Because the fractionation procedure took place under non-denaturing conditions and since aberrant protein aggregation [8] is a common hallmark of both AD and PD, it was hypothesized that macromolecular complexes, potentially altered between the studied groups, could be captured through the HMW fractionation approach.

The present results confirmed this premise, as evidenced by the fact that although no variation was observed in the overall serum protein captured by both strategies, proteins exclusively altered in the HMW fraction accounted for 40% of the total list of altered proteins. Furthermore, with the exception of one protein (Centrosome-associated protein CEP250), all proteins had a MW below 90 kDa, which is considerably lower than the 300 kDa cut-off filter used. It is worth noting that 72% of proteins altered in the HMW fraction did not exhibit alterations in their total levels. This supports the possibility that different regulatory mechanisms of these proteins, apart from expression and degradation, are being revealed and studied using this approach.

Moreover, the results show that most of these proteins have several reported interactors and thus may be involved in the formation of large complexes. An illustration of this phenomenon is found in the protein clusterin (CLU gene), also known as apolipoprotein J, which has been implicated in the metabolism of aggregation-prone proteins, including those associated with NDs [31–33]. Notably, clusterin’s interaction with Aβ42 has been demonstrated to enhance its clearance from the brain through the blood-brain barrier (BBB) [31]. Furthermore, clusterin has been implicated in various stages of PD, potentially exerting a neuroprotective effect through its interaction with α-synuclein aggregates [32]. Additionally, the interaction between clusterin and α-synuclein has been detected in plasma samples [33]. Beyond this specific example, the generic functional analysis of altered proteins, particularly those from the HMW serum strategy, suggests their close association with amyloids and clot formation, which may involve large structures.

On the other hand, some of these proteins may instead, or additionally, be present in large biological structures not composed exclusively of proteins, like exosomes or lipoproteins, which would not only likewise justify their presence in this HMW serum fraction but also give further understanding of the potentially altered mechanisms related to the diseases being studied. Such may be the case of the proteins clusterin and serum amyloid A-4 (*SAA4* gene), which were found to be altered in serum neuron-derived exosomes of AD patients [34]. Additionally, exosomal clusterin was found to be altered in patients at different stages of PD when compared to controls [35]. Thus, although the presence of exosomes in the HMW fraction was not confirmed, given the MWCO of the filters used in this work, it is feasible that some of the proteins being analyzed in the HMW fraction may correspond to proteins linked to the extracellular vesicles.

Altogether, these findings support the notion that the HMW fractionation approach can provide a new level of information that may offer new insights into how proteins are organized within a given sample.

### Altered lipoprotein metabolism can be a peripherical marker of AD and PD

The combination of the two approaches in this study led to a robust and promising potential biomarker panel composed of ten proteins quantified in whole serum or HMW serum. A major finding revealed by this model was the involvement of several lipoproteins in discriminating between the studied groups. Among the ten proteins used in the best discriminant model, three are apolipoproteins: Apo-AI from whole serum, and Apo-CI and Apo-E from HMW serum. Besides those three proteins, other altered apolipoproteins were observed in this study but not included in the model, namely: i) the Apo-AII, Apo-LI and clusterin highlighted in the HMW serum strategy; ii) the Apo-AIV from the whole serum; and, iii) beta-2-glycoprotein 1 in both approaches. Furthermore, a lipoprotein-related enzyme, lecithin-cholesterol acyltransferase, was also altered in HMW serum. This is further supported by the functional enrichment analysis of the altered proteins discovered in both strategies that highlight the involvement of those proteins in lipoprotein metabolism and HDL-mediated lipid transport pathways. Cumulatively, all these findings point to the relevance of lipoproteins in the context of NDs and, although not absolutely clear, the link between these diseases, in particularly AD, and apolipoproteins has been the focus of many studies [31, 33, 36–46].

Apo-AI and Apo-E have an established relation to toxic species clearance from the brain in AD and PD [31, 33, 37, 47–50]. Additionally, regarding AD, our findings for both proteins contradict what can be found in the literature [38, 43]. For Apo-AI, we found an increase in abundance in AD patients as opposed to the decrease reported for most studies [38]. However, in another study where no significant alterations in total serum Apo-AI content of AD patients were reported, further investigation revealed that some proteoforms of this protein were significantly increased compared to the levels observed in the controls [39]. This has been suggested as a possible explanation for the different observations regarding this protein in the context of AD, which might be related to the use of different detection methods within different studies [38]. Regarding this protein’s connection to clearance mechanisms, evidence suggests that for HDLs containing Apo-AI (APOA1-HDL), the structure seems to influence not only the disaggregation of Aβ fibrils but also its ability to cross the BBB, with lipid-poor discoidal APOA1-HDL having the best performance when compared to APOA1-HDL in other lipidation states [37]. Moreover, in the present study, phosphatidylcholine-sterol acyltransferase, an enzyme known to affect HDL structure through lipidation of Apo-AI in plasma [45], was also found to be altered in AD patients. As this enzyme participates in HDL maturation in plasma [51], this result could reflect the dysregulation of lipoproteins in AD. For Apo-E, we found a decrease in abundance in AD patients as opposed to the increase reported for MCI patients in a previous study [43]. Isoform and lipidation status of Apo-E is also crucial for the Aβ clearance, with the Apo-E4 isoform, the genetic risk factor most associated with the onset of AD [31, 48], and higher lipidation having detrimental effects on the process [31, 48]. Further investigation using the combined approach presented in this study, particularly in the context of AD, should also involve a lipid profile analysis and APOE genotyping of the participants to enable a more comprehensive characterization of serum HDLs and Apo-E content, respectively.

Although systemic lipid abnormalities have also been implicated in PD, there are fewer findings connecting it to HDL-related proteins, as compared to AD [32]. Nonetheless, a previous report observed significantly decreased values of Apo-AI in mild PD patients compared to healthy controls, but much like what we observed for this protein, a less impactful and non-significant decrease was observed in moderate/severe PD patients [52]. In fact, most research indicates that Apo-AI may have a protective role in PD [32] and it has been hypothesized that APOA1-HDL could take part in the efflux of α-synuclein from the brain [33]. Additionally, both Apo-AI and Apo-E have been reported to interact with α- synuclein [33].

Finally, HDL size and plasma levels have been shown to be dependent on the levels of Apo-CI [53], another apolipoprotein included in the best diagnostic model generated in this study. Besides that, a previous study also shows that the dysregulation of this protein can lead to impaired memory processes in mice [54]. This suggests that the regulation of Apo-CI can be pivotal in the brain and that a systemic disruption of this process could have effects detectable beyond the CNS, particularly in lipoprotein metabolism, that could be observable in peripheral biofluids. Interestingly, this study found that Apo-CI significantly decreased in the HMW serum fraction of AD compared to PD patients, but only a slight and non-significant decrease was observed when compared to CT patients. Similarly, only a small non- significant decrease in HMW serum Apo-CI was observed for controls in comparison to PD patients, which is in accordance to what was already reported for whole plasma [40]. Given that memory impairment is a hallmark of AD and not a predominant feature among PD patients [55], these results may be influenced, at least in part, by the age disparities observed between the PD group and the other two groups. In fact, PD patients are, on average, younger than those in the other two groups, with similar age distributions. Additionally, the results regarding Apo-CI may also be affected by individual Apo-CI and Apo-E isoforms. Notably, the *APOE* and *APOC1* genes are in linkage disequilibrium [56], and carriers of the *APOE(ε4)* and *APOC1(H2)* alleles have been demonstrated to have an increased risk of developing AD [46]. This was further confirmed in a study using human *APOE*-carrying mice, which demonstrated that those animals carrying the *APOE(ε4)* allele were found to have decreased serum Apo-CI content when compared to those carrying the *APOE(ε3)* allele [57]. However, Apo-CI has also been suggested to potentially play a modulatory role in the development of AD, with reported effects on mice cognitive function independent of Apo-E expression [58]. Again, these observations strengthen the importance of combining these results with further characterization of the individuals, including genotyping of the apolipoproteins’ isoforms.

Although these three apolipoproteins have been linked to AD and PD, as evidenced by previously mentioned studies, and their relevance for the discriminant model, their use alone or in combination with other proteins associated with apolipoprotein-related mechanisms did not result in a robust diagnostic model capable of effectively distinguishing between the studied groups (as shown in Additional File 1: Supplementary Figure 7). This indicates that these three apolipoproteins had to be combined with other seemingly unrelated proteins to be used as potential biomarkers. Further studies should be directed towards elucidating this potential relationship to understand: i) the importance of the identified proteins/mechanisms for the pathophysiology of the studied NDs, and ii) to which extent these mechanisms are differently altered between the two diseases.

In summary, this work demonstrated that combining two complementary sample processing approaches is a more effective strategy to reach potential biomarkers than a single approach. Besides that, the strategies used here, which combine the analysis of the whole serum and the HMW fractionation of non- denaturing serum, can also identify proteins being differentially modulated besides the conventional alteration in their total levels. In this work, this new strategy was applied to a cohort of NDs patients and respective CT individuals, being able to build a good predictive model capable of distinguishing all the three groups studied (AD, PD and CT). This predictive model highlighted the linkage of the apolipoprotein family and NDs, with three out of the ten proteins included in this model being apolipoproteins. Nevertheless, further validation in a larger and independent cohort is needed to confirm the robustness and reliability of the model, as well as more studies to link the alterations observed and these pathologies. Controlling the lipid profile in future studies is also advised, as altered lipid metabolism was a major finding of the present work. Another aspect to be further explored would be the identification of protein complexes in the HMW fraction to better understand the origin of the protein alterations observed.

Overall, the results of this study demonstrate that HMW fractionation under non-denaturing conditions could be a valuable addition to routine biofluid analysis, particularly regarding the detection of changes in macromolecular organization. Furthermore, this study highlights the importance of exploring the involvement of apolipoproteins and lipid metabolism in greater detail within the context of NDs.

## abbreviations

AD: Alzheimer’s Disease;
APOA1-HDL: High-Density Lipoproteins containing Apolipoprotein A1;
AUC: Area Under the Curve;
Aβ: Amyloid-β;
BBB: Blood-Brain Barrier;
BioGRID: Biological General Repository for Interaction Datasets;
CE: Collision Energy;
CES: Collision Energy Spread;
CHUC: Centro Hospitalar e Universitário de Coimbra;
CI: Confidence Interval;
CNS: Central Nervous System;
CT: Healthy Controls;
DDA: Data-dependent Acquisition;
DMSO: Dimethyl Sulfoxide;
FA: Formic Acid;
FDR: False Discovery Rate;
GO: Gene Ontology;
HDL: High-Density Lipoproteins;
HMW: High Molecular Weight;
IS: Internal Standard;
LDA: Linear Discriminant Analysis;
MCI: Mild Cognitive Impairment;
MS: Mass Spectrometry;
MW: Molecular Weight;
MWCO: Molecular Weight Cut-Off;
NDs: Neurodegenerative Diseases;
PD: Parkinson’s Disease;
ROC: Receiver Operating Curve;
SE: Standard Error;
STRING: Search Tool for Retrieval of Interacting Genes/Proteins;
XIC: Extracted Ion Chromatogram.

## Acknowledgements

All authors have read the journal’s authorship statement. The authors would like to thank the financial support of the European Regional Development Fund (ERDF), through the COMPETE 2020 - Operational Programme for Competitiveness and Internationalisation and Portuguese national funds via FCT – Fundação para a Ciência e a Tecnologia, I.P., under projects: MEDPersyst - POCI-01-0145-FEDER-016428 (ref.: SAICTPAC/0010/2015), POCI-01-0145-FEDER-30943 (ref.: PTDC/MEC-PSQ/30943/2017), PTDC/MED-NEU/27946/2017, POCI-01-0145-FEDER-016795 (ref.: PTDC/NEU-SCC/7051/2014), POCI-01- 0145-FEDER-029311 (ref.: PTDC/BTM-TEC/29311/2017); EXPL/BTM-TEC/1407/2021, and, UIDB/04539/2020, UIDP/04539/2020, UIDB&P/4950/2020 and LA/P/0058/2020, and the National Mass Spectrometry Network (RNEM) [POCI-01-0145-FEDER-402-022125 Ref. ROTEIRO/0028/2013). SIA was supported by the CEEC grant 2021.04378.CEECIND and MR was supported by Ph.D. fellowship 2020.07749.BD.

## Ethics approval

The study was approved by the Ethics Committee of the Faculty of Medicine of the University of Coimbra (reference CE_010.2017) and the Ethics Committee of the Centro Hospitalar e Universitário de Coimbra (CHUC) (reference 34 /CES-CHUC-024-18) and was conducted according to the principles stated in the Declaration of Helsinki [59]. Written informed consent was obtained from all participants.

## Data linking

The MS proteomics data have been deposited to the ProteomeXchange Consortium [60] via the PRIDE [61] partner repository with the dataset identifier PXD034077 (for reviewer: Username; Password: jhF8zQaR).

## Authors’ contributions

SIA and BM conceived and designed the study. SIA and MR performed all the experiments, data analysis, and wrote the manuscript. IB, AG, DP, CS, JP, CJ, IS, AV, AM, and MCB were involved in the selection and collection of the plasma samples used in this study. MCB and BM were responsible for funding. The author(s) read and approved the final manuscript.

