## Additional File 1 for "Unmasking Hidden Systemic Effects of Neurodegenerative Diseases: A Two-Pronged Approach to Biomarker Discovery"

Provide the full postal address of each affiliation

^1^ CNC - Centro de Neurociências e Biologia Celular, Universidade de Coimbra, Rua Larga - Faculdade de Medicina, 1ºandar - POLO I, 3004-504 Coimbra, Portugal.

^2^ CIBB - Centre for Innovative Biomedicine and Biotechnology, University of Coimbra, Rua Larga - Faculdade de Medicina, 1ºandar - POLO I, 3004-504 Coimbra, Portugal.

^3^ Instituto de Investigação Interdisciplinar, Universidade de Coimbra (IIIUC), Casa Costa Alemão, R. Dom Francisco de Lemos, 3030-789 Coimbra, Portugal.

^4^ Faculdade de Medicina, Universidade de Coimbra, Azinhaga de Santa Comba, 3000-548 Coimbra, Portugal.

^5^ - University of Coimbra, Institute for Interdisciplinary Research, Doctoral Programme in Experimental Biology and Biomedicine (PDBEB), Portugal

^6^ Serviço de Neurologia, Centro Hospitalar e Universitário de Coimbra, Praceta Professor Mota Pinto, 3004-561 Coimbra, Portugal.

^7^ Coimbra Institute for Biomedical Imaging and Translational Research (CIBIT), Azinhaga de. Santa Comba, 3000-548 Coimbra, Portugal.

^8^ Department of Neurosciences and Mental Health, Santa Maria Hospital - CHLN, ISAMB, University of Lisbon, Av. Prof. Egas Moniz MB, 1649-028 Lisboa, Portugal.

^9^ Instituto de Medicina Molecular e Faculdade de Medicina, Universidade de Lisboa, Av. Prof. Egas Moniz, 1649-028 Lisboa, Portugal.

* - Equal contribution

\# - Senior equal contribution

† - Corresponding authors: Bruno Manadas (, +351231249170) and Sandra Anjo (, +351231249170), Center for Neuroscience and Cell Biology – University of Coimbra, UC Biotech - Parque Tecnológico de Cantanhede, Núcleo 04, Lote 8, 3060-197 Cantanhede – Portugal

**Supplementary Figures**

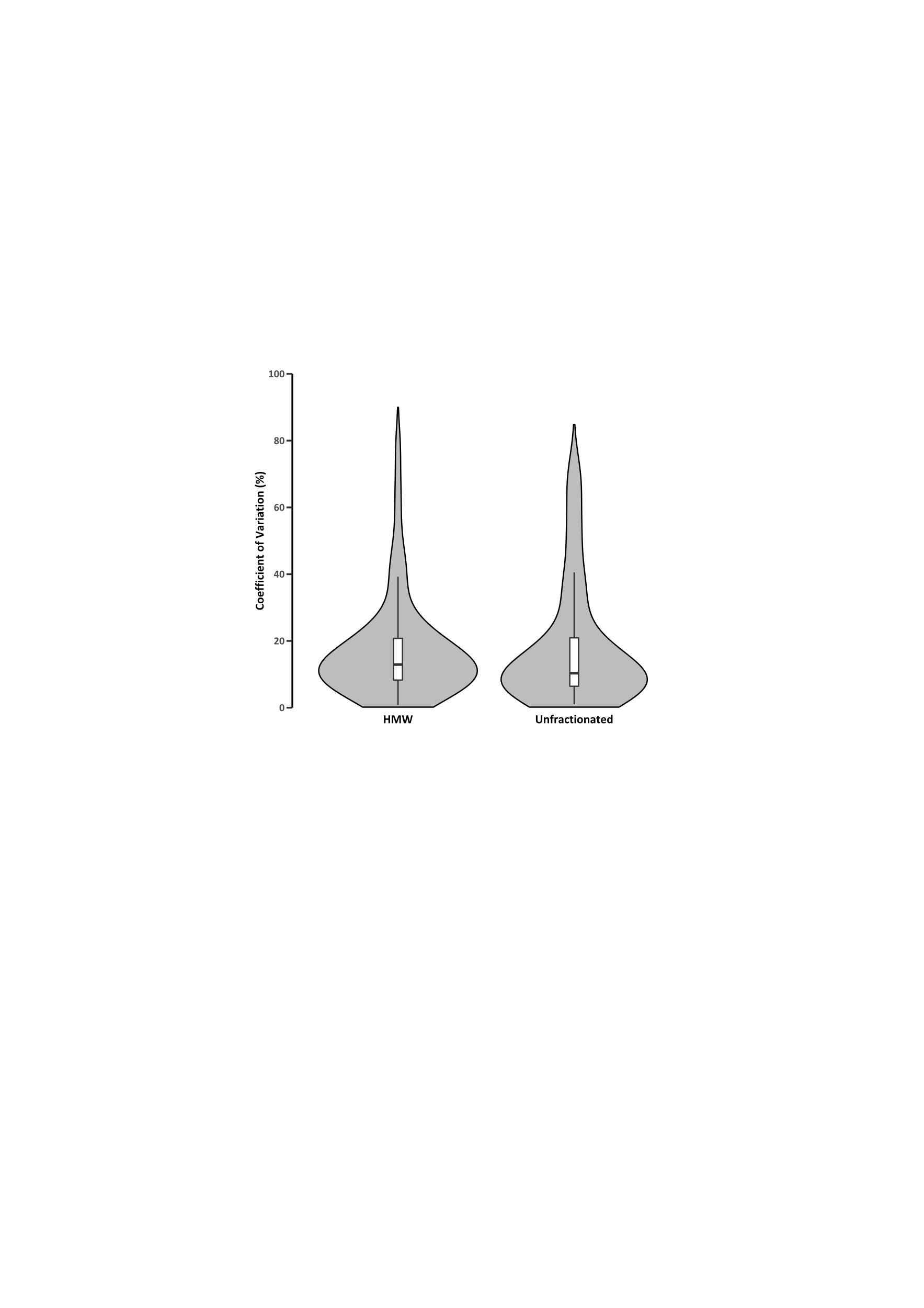

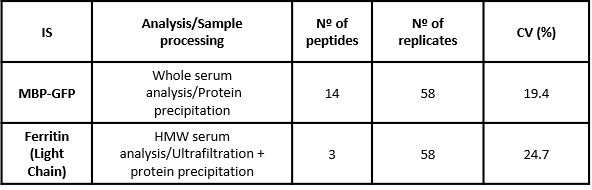

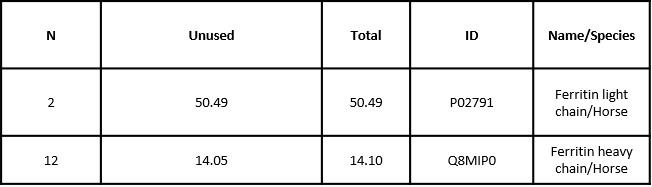

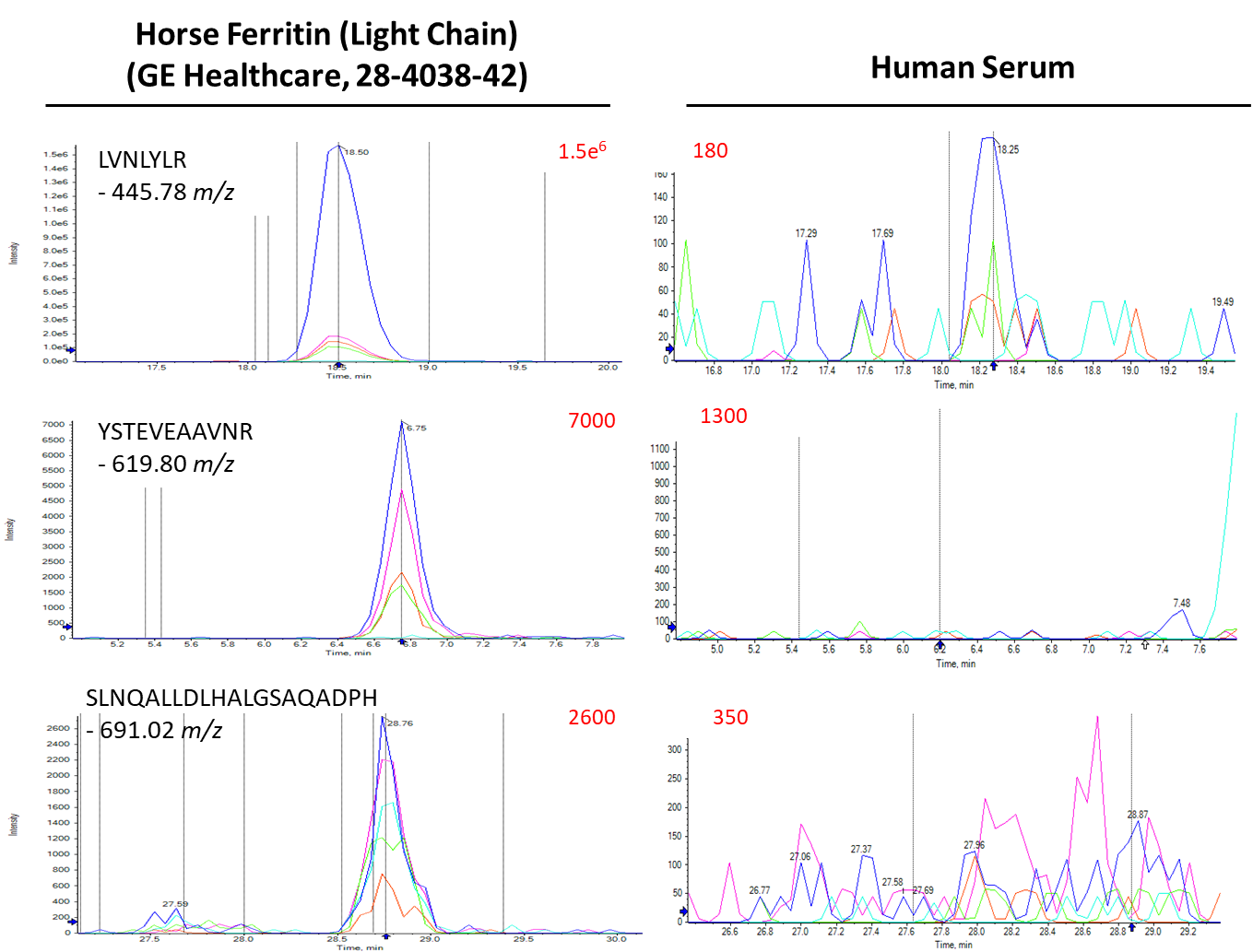

**(a)**

**(d)**

**(b)**

**(c)**

**Supplementary Figure 1:** **Internal standard proteins and ultrafiltration reproducibility.** (a) Identification of HMW internal standard protein (Equine Ferritin) spiked in serum samples. (b) Serum matrix effect on HMW internal standard protein quantification. (c) Variability of quantification values, acquired for internal standard proteins used in each sample preparation approach. (d) Variability distribution of quantified proteins in HMW serum and (unfractionated) whole serum sample replicates. Non-normalized data from 3 replicates of each approach was used, comprising 129 proteins quantified in HMW serum and 174 proteins quantified in whole serum. In both approaches 75% of quantified proteins had a coefficient of variation below 21%. HMW – High Molecular Weight. (1)

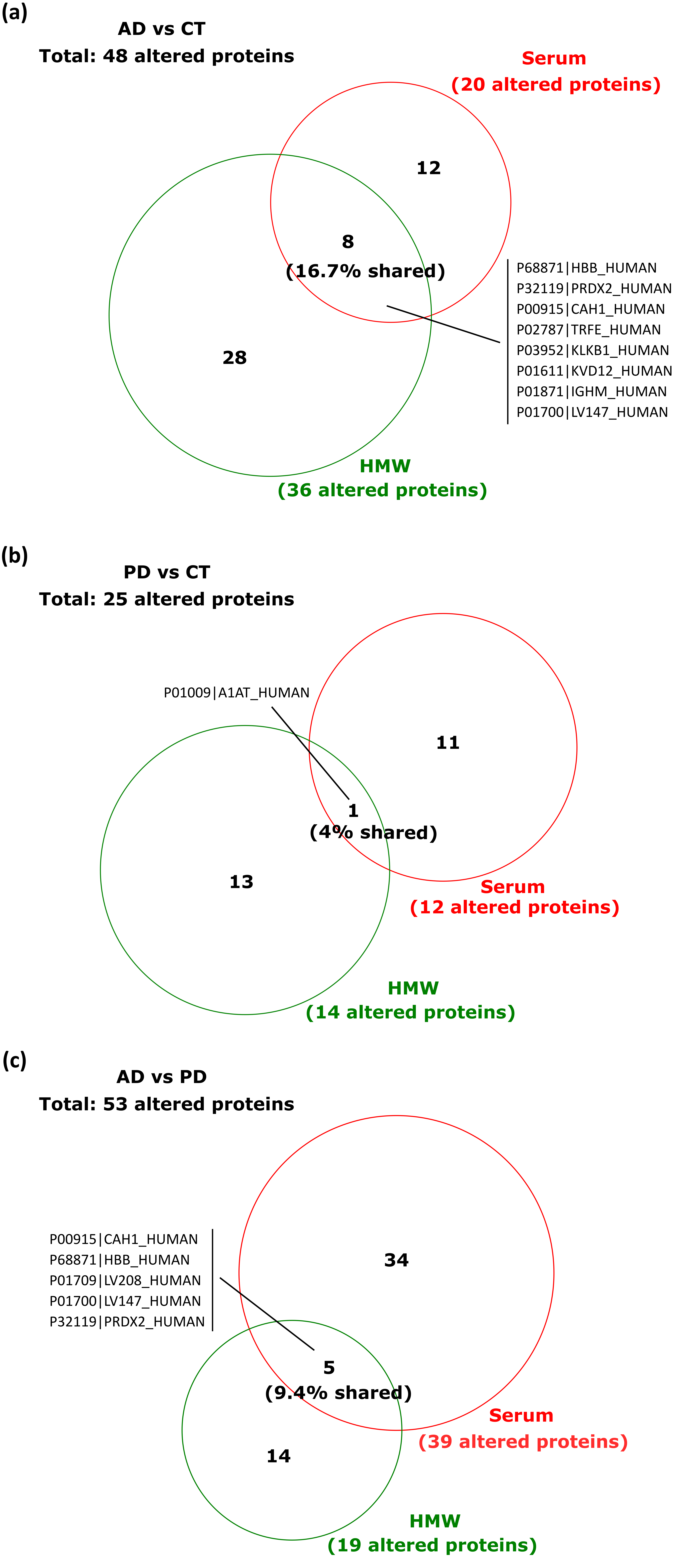

**Supplementary Figure 2: Altered proteins quantified using the whole serum dataset and HMW serum dataset.** (a) Venn diagram comparing the number of quantified proteins altered with statistical significance between AD and CT using the whole serum dataset (red) and the HMW serum dataset (green). (b) Venn diagram comparing the number of quantified proteins altered with statistical significance between PD and CT using the whole serum dataset (red) and the HMW serum dataset (green). (c) Venn diagram comparing the number of quantified proteins altered with statistical significance between AD and PD using the whole serum dataset (red) and the HMW serum dataset (green). Statistical analysis was performed by a Kruskal–Wallis H test followed by the Dunn's Test for pairwise comparison. Dunn's p-value were corrected using the using the Benjamini-Hochberg FDR adjustment and the statistical significance was considered for p-values below 0.05. Shared altered protein identifiers are highlighted. HMW – High Molecular Weight; AD – Alzheimer’s Disease; PD – Parkinson’s Disease; CT – healthy controls.

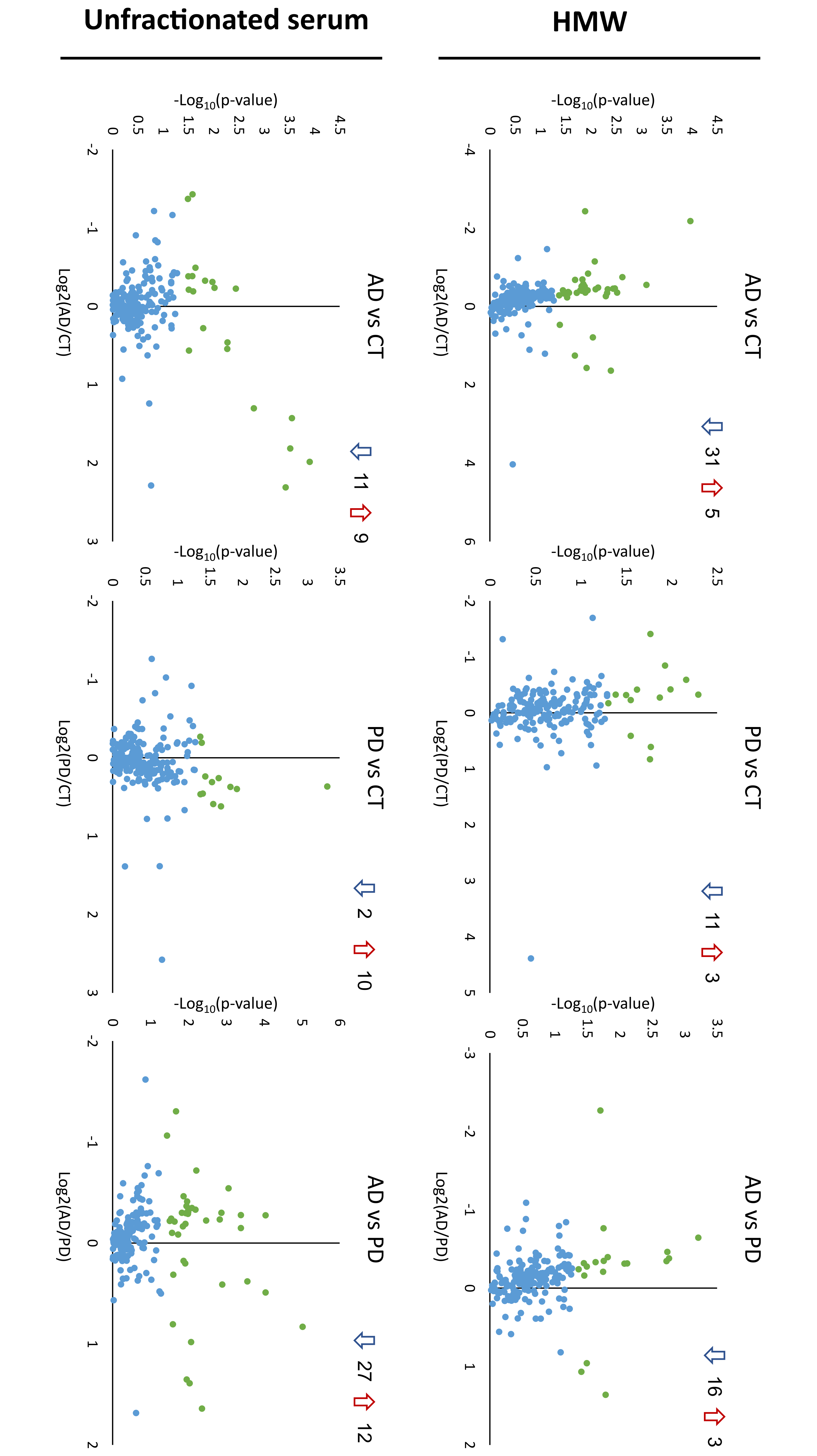

**Supplementary Figure 3: Volcano plots comparing all proteins quantified across the three different experimental groups using the two different serum-processing strategies.** Statistical analysis was performed by a Kruskal–Wallis H test followed by the Dunn's Test for pairwise comparison. Dunn's p-values were corrected using the using the Benjamini-Hochberg FDR adjustment and the statistical significance was considered for p-values below 0.05. Green dots represent proteins altered with statistical significance. In each pairwise group comparison, blue downward arrows indicate the number of altered downregulated proteins and red upward arrows indicate the number of altered upregulated proteins. The first row represents data from the HMW serum dataset and the second row represents data from the whole serum dataset.

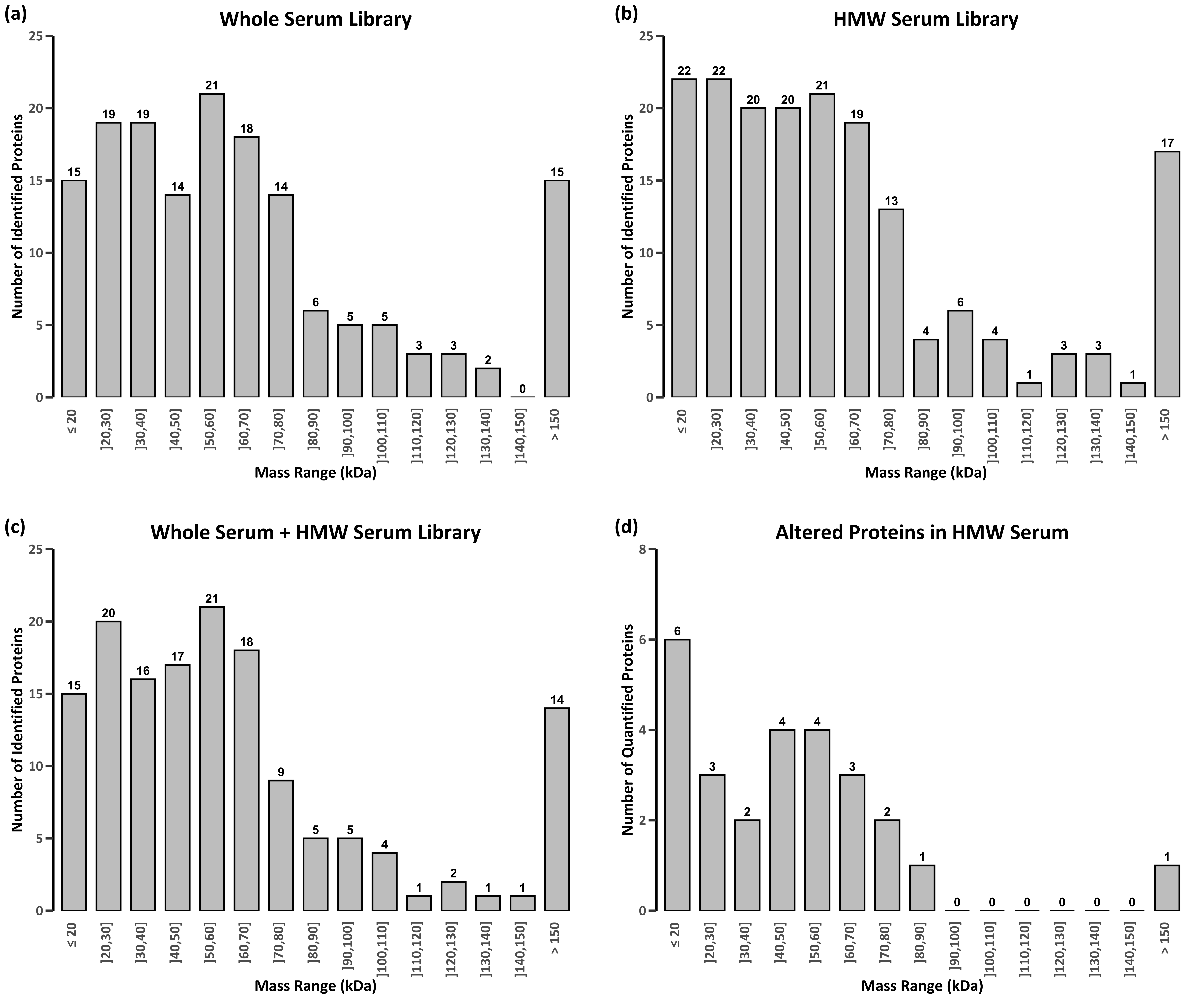

**Supplementary Figure 4: Mass distribution of proteins identified or quantified using different approaches.** (a) Molecular weight distribution of proteins identified using the whole serum approach; (b) Molecular weight distribution of proteins identified using the serum ultrafiltration approach (MWCO ≥ 300 kDa); (c) Molecular weight distribution of proteins identified by searching data from both approaches (whole serum and HMW serum); (d) Molecular weight distribution of significantly altered proteins (p-value below 0.05 according to Kruskal–Wallis H test, with Dunn's Test correction and Benjamini-Hochberg adjustment) quantified only by using the serum ultrafiltration approach. Immunoglobulins are omitted. MWCO – Molecular weight cut-off; HMW – High Molecular Weight.

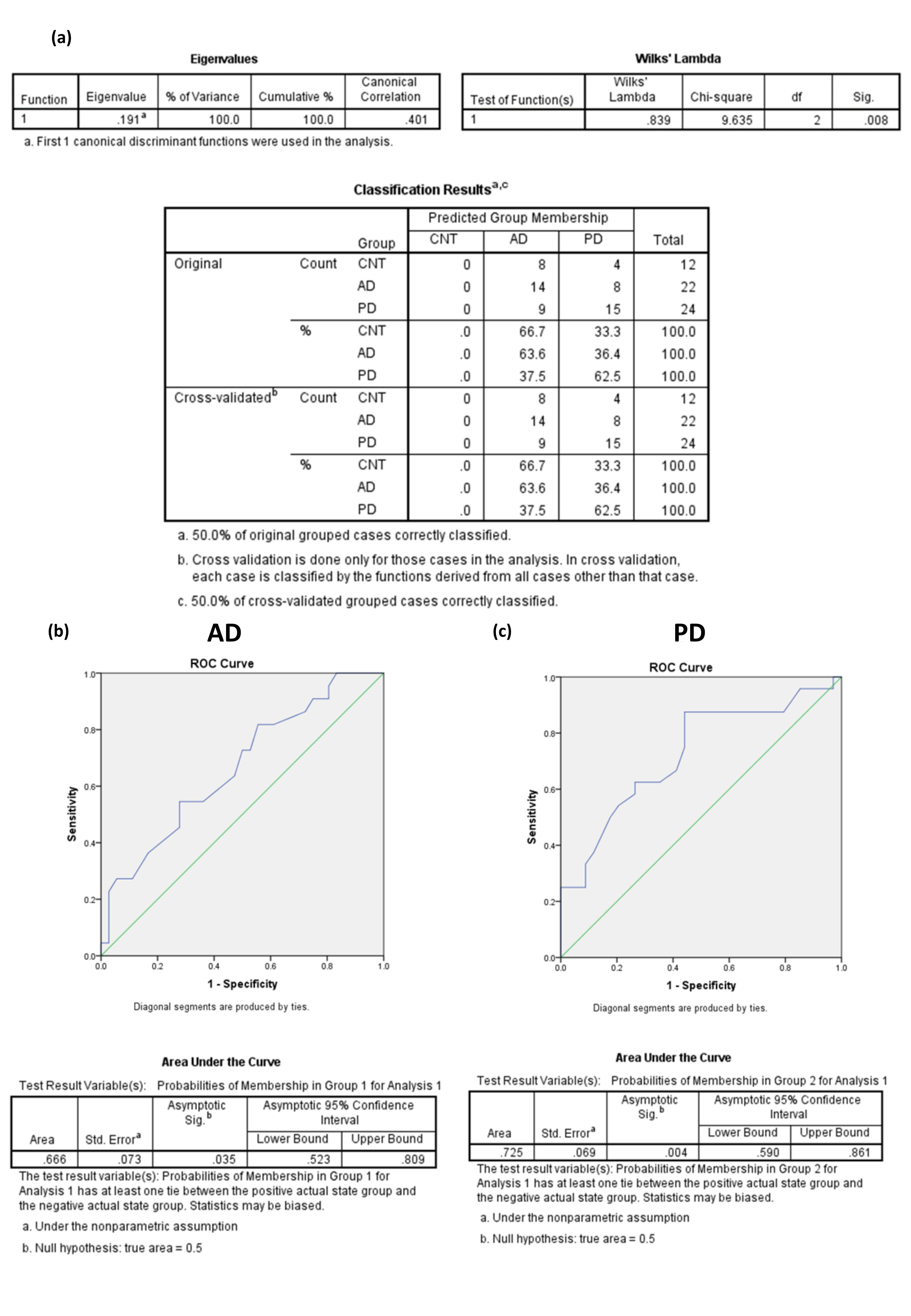

**Supplementary Figure 5: Linear discriminant analysis using patient’s age.** (a) Summary statistics and classification of samples according to a discriminant model built using patient’s age. (b) Comparative ROC curve of the discriminant function generated for AD samples and the respective AUC and 95% CI. (c) Comparative ROC curve of the discriminant function generated for PD samples and the respective AUC and 95% CI. ROC - Receiver operating characteristic; AUC - Area under the curve; CI – Confidence interval; AD – Alzheimer’s Disease; PD – Parkinson’s Disease.

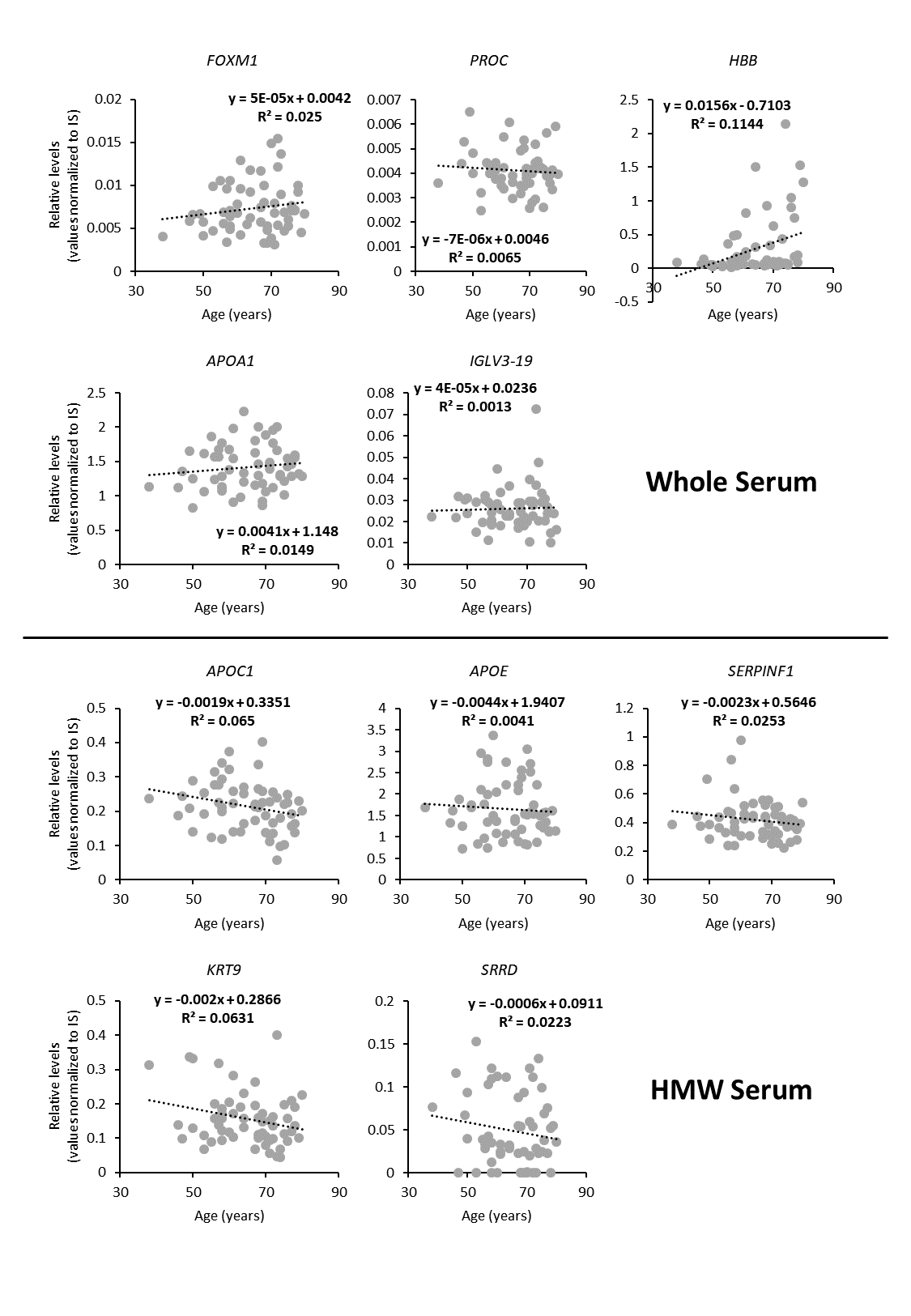

**Supplementary Figure 6: Correlation between quantification of the 10 proteins included in the combined model and patient age.** Upper panel shows proteins selected from the whole serum approach and lower panel shows proteins selected from the HMW serum approach. R^2^ - coefficient of determination.

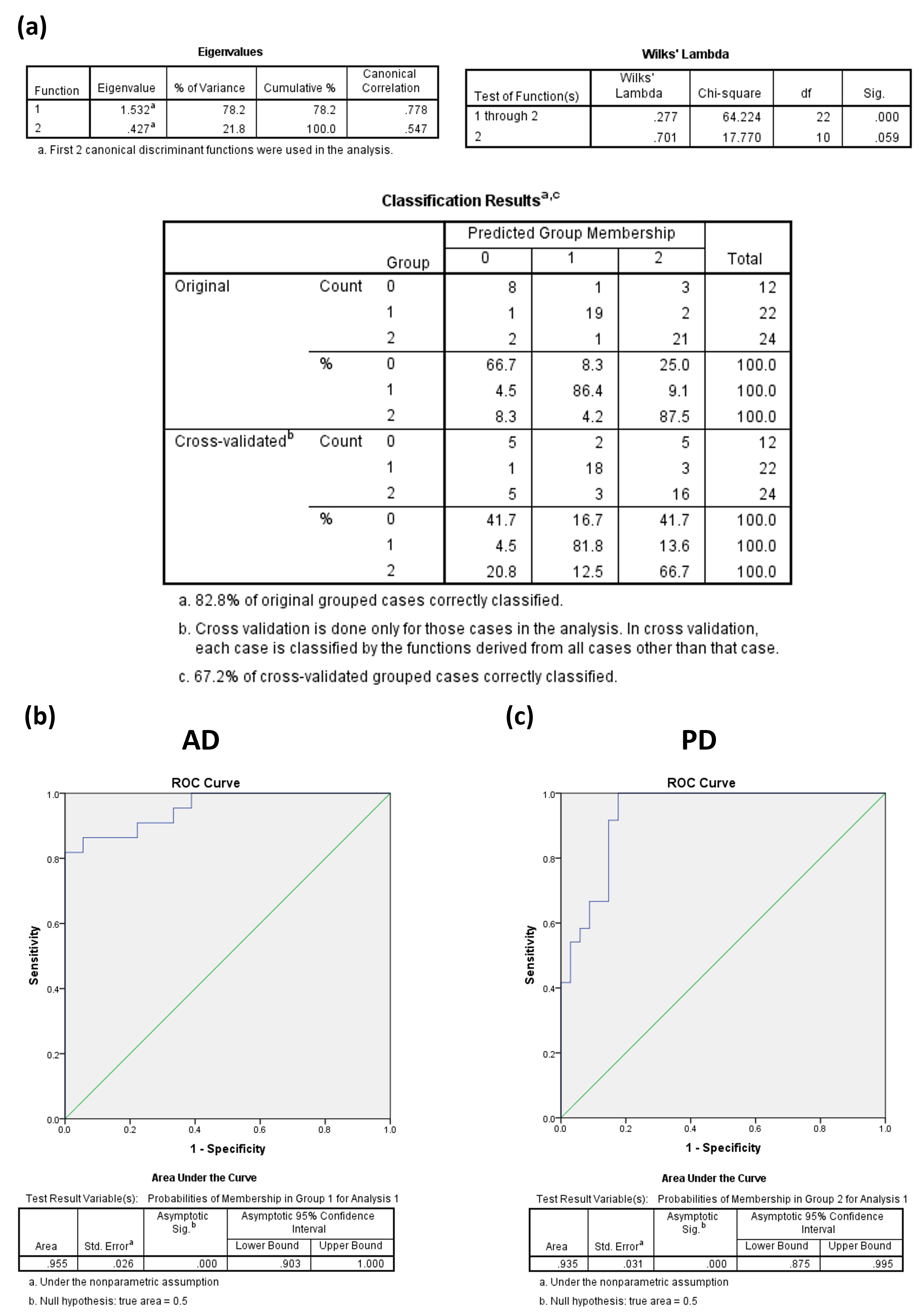

**Supplementary Figure 7: Linear discriminant analysis using 10 altered proteins associated with apolipoprotein related mechanisms.** (a) Summary statistics and classification of samples according to a discriminant model built using data from 10 altered proteins involved in apolipoprotein related mechanism – Apo A2, Apo C1, Apo E, Apo L1, Clusterin, Beta-2-glycoprotein and LCAT enzyme from the HMW serum dataset and Apo A1, Apo A4, Albumin and Beta-2-glycoprotein from the whole serum dataset. (b) Comparative ROC curve of the discriminant function generated for AD samples and the respective AUC and 95% CI. (c) Comparative ROC curve of the discriminant function generated for PD samples and the respective AUC and 95% CI. HMW – High Molecular Weight; ROC - Receiver operating characteristic; AUC - Area under the curve; CI – Confidence interval; AD – Alzheimer’s Disease; PD – Parkinson’s Disease.

**Supplementary Tables**

**Supplementary Table 8 – Summary of the specificity and sensitivity of the LDA methods obtained using all the altered proteins per sample type (whole serum or HMW serum) or the combination of the two (Whole + HMW serum).** Values are in percentage.

|  | **Specificity** | **Sensitivity** | |
| --- | --- | --- | --- |
| **Model** |  | **AD** | **PD** |
| **Whole Serum** | 66.7 | 86.4 | 87.5 |
| **HMW Serum** | 75.0 | 77.3 | 83.3 |
| **Whole + HMW Serum** | 91.7 | 95.5 | 100 |
